# Boron alters carboxyl group binding capacity and Al transport pathway to relieve Al toxicity

**DOI:** 10.1101/2019.12.12.874412

**Authors:** Lei Yan, Muhammad Riaz, Jiayou Liu, Yalin Liu, Yu Zeng, Cuncang Jiang

**Affiliations:** Microelement Research Center, College of Resources and Environment, Huazhong Agricultural University, Wuhan, Hubei, 430070, P. R. China; College of Natural Resources and Environment, South China Agricultural University, Guangzhou 510642, P. R. China

**Author notes:** To whom correspondence should be addressed. Correspondence: Cuncang Jiang, Microelement Research Center, Huazhong Agricultural University, Wuhan, Hubei Province, 430070, P.R. China. **Author contributions** C.C.J. and L.Y. designed and supervised this study. L.Y. conducted the experiments, performed data interpretation, and drafted the manuscript. R.Muhammad and J.Y.L. helped in interpreting the results of the study, and Y.L.L., Y.Z. helped in replacing the nutrition solution in the experiment and determining B and Al concentration using the curcumin colorimetry and inductively coupled plasma-mass spectrometry methods respectively. R.Muhammad helped to revise the manuscript grammatically. All authors read and approved the final manuscript. Author email addresses: LY, MR, JL, YL, YZ.

**Keywords:** Aluminum toxicity, boron, cytoplasm, pectin, trifoliate orange rootstock, vacuoles

## Abstract

Boron (B) is indispensable for plant growth and has been reported in the mitigation of aluminum (Al) toxicity in different plants. This study unraveled the efficacy of B in reducing the toxicity of Al to trifoliate orange seedlings in a hydroponic experiment. In the current study, B supply had a positive effect on root length and plant growth-related parameters, and attenuated Al-induced inhibition of plasma membrane H^+^-ATPase activity. The results of XPS and SEM-EDS revealed that B reduces the Al accumulation in root cell wall (CW), especially acts on pectin fractions (alkali-soluble pectin), accompanied by suppressing the pectin synthesis, inhibiting pectin methylesterase (PME) activity and *PME* expression. Furthermore, B application inhibits *NRAT1* expression while increases *ALS1* expression, which are responsible for restraining Al transport from external cells to the cytoplasm and accelerating Al divert to vacuoles, and the results can be further demonstrated by TEM-EDS analysis. Taken together, our results indicated that B mainly promotes the efflux of H^+^ by regulating the plasma membrane H^+^-ATPase activity, futhur reduce the demethylation of pectin to weaken Al binding ability to carboxyl. More importantly, B alleviated some of the toxic effects of Al by decreasing the deposition of Al in cytoplasm and compartmentalizes Al into vacuoles.

**One-sentence summary:** Boron can reduce the binding amount of carboxyl group to Al in pectin, decreasing the deposition of Al in cytoplasm and compartmentalizes Al into vacuoles, thereby reduce the toxicity of Al to plants..

## 1. Introduction

Aluminum (Al) is the 3rd most abundant element in the earth’s crust, and widely distributed, accounting for approximately 40% of the world’s arable land, and usually exists in the form of non-toxic silicates or other sediments (Kochian, 1995). However, Al solubility enhances and solubilizes into Al^3+^ in acidic soils (pH ≤ 5), which hinders the acquisition of nutrients as a result of inhibition of plant root elongation (Horst *et al*., 2010). Aluminum toxicity impedes root growth by increasing the reactive oxygen species (ROS) generation and destroying the homeostasis of antioxidant metabolism (Mora *et al*., 2016). The initial symptom of Al toxicity is on root growth, leading to changes in root morphology, such as atrophy of root hair, and swelling of root tips (Huang *et al*., 2014) and decreasing cell wall (CW) elasticity and plasticity (Ma *et al*., 2004; Horst *et al*., 2010). In recent years, studies proposed that Al binding to the CW is a prerequisite for Al toxicity to plants (Horst *et al*., 2010; Kopittke *et al*., 2015), and root CW is the chief site of Al^3+^ accumulation and the main barrier to Al toxicity (Horst *et al*., 1995). Plant CW is mainly composed of pectin, cellulose, and hemicellulose. Pectin, as the main component of CW, accounts for about 35% of CW weight in dicotyledons and monocotyledons of non-gramineae, and is mainly concentrated in the primary CW and the mesohyl layer (Smith & Harris, 1999), and is an important platform in Al perception and response (Horst *et al*., 2010). The adsorption and fixation ability of Al to CW depends on the pectin content and the degree of methylation (DM), which together determines the total negative charge carried by pectin. Pectin methyl esterification is regulated by pectin methylesterase (PME) (Schmohl & Horst, 2000), which mainly contributes to the hydrolysis of the carboxyl group of pectin methyl esterification and reduces the degree of methyl esterification by demethylation (Grsic-Rausch & Rausch, 2004). It is widely accepted that PME activity determines the toxicity of Al in plants (Rangel *et al*., 2009). In particular, the negative charges on pectin matrix such as the free carboxyl groups of pectin are the fundamental adsorption sites of Al^3+^ (Yang *et al*., 2011a; Yan *et al*., 2018a). It has been proposed theoretically that hemicellulose also plays an important role in the adsorption of Al, it is also the main Al^3+^ binding site in CW (Yang *et al*., 2011a). Hemicellulose 1 (HC1) is the largest accumulator for Al^3+^ in CW (Yang *et al*., 2011a; Yang *et al*., 2011b). It has been reported that more than 85% and 62% of the Al in CW accumulated in the HC1 of the Al-sensitive and Al-resistant alfalfa varieties, respectively (Wang *et al*., 2017). Plants have developed a variety of detoxification strategies to cope with Al toxicity. These include shielding the CW of Al-binding site (Yan *et al.*, 2018a), reducing the transfer of Al from the external environment to cells and detoxifying Al inside the apoplast (Zhu *et al*., 2018), and compartmentalizes Al into the vacuoles (Xia *et al*., 2010). Therefore, enhancing the exclusion of Al from root CW is the pivotal role of plants to resist Al toxicity (Zhu *et al*., 2019).

Boron (B) is an essential microelement for normal plant development (O’Neill *et al*., 2001) and is proposed in many plants to reduce the toxicity of Al. Boron supply to Al-containing hydroponic nutrient media increases root elongation in higher plants such as *Citrus grandis* (Zhou *et al*. 2015), pea (Li *et al*. 2017), rape seedlings (Yan *et al*. 2018b), and trifoliate orange (Riaz *et al*., 2018). Boron can alleviate Al toxicity by reducing the Al^3+^ binding sites (carboxyl groups) in CW. Moreover, increased B application reduced root oxidative damage by changing CW composition and structure (Yan *et al*., 2018a). Another study found that B affects pectin content and the DM of pectin to determine the CW’s ability to adsorption and desorption of Al (Horst *et al*., 2010; Yan *et al*., 2018a). Indeed, the mitigation mechanisms of B to Al toxicity in plants has been studied. Zhu *et al*., (2019) pointed out that B alleviates Al toxicity by altering pectin content and its properties. Moreover, B could enhance Al transfer from the cytoplasm to vacuoles by regulating Al-related transporters in rice root (Zhu *et al*., 2019). However, the mechanisms of B reducing Al toxicity in different plants may be distinct and complex.

Citrus is a fruit crop globally cultivated for human consumption as a main source of carotene, vitamins, carbohydrate, minerals, and other nutritional constituents. Ganzhou, Jiangxi province is one of the suitable regions for citrus plantation with abundant citrus resources and long plantation history, and the production of citrus could be rated the first level in the world (Deng, 2014). However, in this area, 82.1% of the acid red soil is accompanied by B-deficiency due to the high precipitation (Wang *et al*. 2011). Therefore, both Al toxicity and B-deficiency inhibit the yield and quality of citrus (Jiang *et al*., 2009; Shah *et al*., 2017). Trifoliate orange (*Poncirus trifoliate* (L.) Raf.), is a world-famous citrus rootstock and sensitive to B-deficiency (Liu *et al*., 2013). Our previous study has shown that B application on the Al-stressed citrus improved the root elongation by altering CW components and structure (Yan *et al*., 2018a). However, there is no specific study on how B alleviates Al toxicity by affecting CW components and Al transfer in root tip cells, especially focused on the physiology and molecular reactions of Al in trifoliate orange rootstocks. The aims of our study were to investigate the components of the CW which are involved in the alleviation of Al toxicity by B supply and its underlying mechanisms (1), exploring a new insight into the mechanisms of B regulating Al transport in root tips (2). This is the first study to devoting the research of different components of CW fractions in citrus roots after different B levels under Al toxicity, and combined with the actual growth environment, providing a theoretical basis for the application of B-containing fertilizers to acid soils to reduce the Al toxicity in citrus.

## 2. Material and methods

### 2.1. Plant material and experimental conditions

Healthy and pathogens free seeds of trifoliate orange (*Poncirus trifoliate* L. Raf.) were germinated as described previously by Yan *et al*., (2019). After germination, the seeds were cultured in a 1/16 strength Hoagland & Arnon, (1950) nutrient solution until the seedlings grew to the roots about 5.8 cm long, and then uniform seedlings were transferred to 2 L pots (4 seedlings each pot) with 1/4 strength Hoagland & Arnon, (1950) nutrient solution (pH=4.5) containing 0.1, 5, 10, or 50 µM B with or without 300 μM Al (AlCl_3_·6H_2_O). All seedlings were administered to the following eight groups (–Al B0.1 (CK), –Al B5, –Al B10, –Al B50, +Al B0.1, +Al B5, +Al B10, +Al B50). The Al concentrations in the solution were defined in preliminary studies. According to the ionic activity formula and Debye-Hückel equation (Manov *et al*., 2002), the actual chemical activity of Al^3+^ in nutrient solution was 201.09 µM. Seedlings were grown in a greenhouse at 25 °C and photoperiod of 16/8 h day/night at Huazhong Agricultural University (HZAU), Wuhan, China. Replacement of nutrient solution was carried out every after 7 days and first acclimatized in 1/4 strength, and then 1/2 strength, finally at the full strength, continued for 90 days in total and were constantly aerated for 20 min every 4 h. Every set of experiments consisted of three replicates.

Similarly, in order to explore the transfer and accumulation of Al in different organelles of root tip, we further performed a hydroponic experiment, the trifoliate orange rootstocks seedling roots with 6.9 cm were grown by the same seedling-raising method, and then they were transplanted to the hydroponic system with Hoagland & Arnon’s nutrient solution at the same light intensity and temperature as described above. There were two Al treatments (0 μM, –Al; 300 μM, +Al) as well as three B treatments (0.1 µM, B0.1; 10 µM, B10; 50 µM, B50) in the nutrient solution system. After 60 days, the same growth indicators of seedlings as those in the previous hydroponic experiment were detected and are shown in Fig. S1. Boron and Al content are shown in Table S1.

### 2.2. Measurement of growth

The seedlings cultured in nutrient solution were respectively taken out and then separated into roots and shoots, and washed separately with double distilled water and dried on paper towels, and then weighed with an electronic analytical balance. The plant height and root length were measured by a graduated ruler. The roots and shoots from one plant per experimental unit were dried to constant weight in a forced ventilation oven 75 °C. The dry weight was measured and ground. Other samples were frozen in liquid N_2_ immediately after being removed from the nutrient solution and then stored at -80 °C until the extraction of enzymes and CW.

### 2.3. Analysis of total B and Al content in root and Al content in root tips

The 0.2 g of dried root samples were carbonized in an electric furnace, then ashed in a muffle furnace at 500 °C for 4 h. And 10 mL of 0.1 M HCl was added to these samples after cooling at room temperature and the filtrate was collected for further testing. Boron content was measured using Dible *et al*., (1954) method.

The total Al content in the filtrate was determined by inductively coupled plasma-mass spectrometry (ICP-OES) (Agilent Technologies 5110, USA) (Yu *et al*., 2009). For root tips Al determination, the root segments (0-10 mm, 10 root tips for one replicate) were extracted with 2 M HCl for 24 h, shaking continuously during the extraction, and then determined by ICP-OES (Yu *et al*., 2009).

### 2.4. Cell wall (CW) extraction and fractionation

The root CW was isolated according to Hu & Brown, (1994) method. Root CW materials (pectin, hemicellulose 1 (HC1), and hemicellulose 2 (HC2)) were employed by Zhong & Läuchli, (1993) method. Three types of pectin (water-soluble pectin (WSP), chelator-soluble pectin (CSP), and alkali-soluble pectin (ASP)) were extracted by the Redgwell & Selvendran, (1986) method (detail can be found in the supplementary data).

### 2.5. Measurement of the contents of pectin, hemicellulose 1, and hemicellulose 2

The uronic acid content in pectin was measured colourimetrically at a wavelength of 520 nm by Blumenkrantz & Asboe-Hansen, (1973) method with galacturonic acid as the standard. The content of total sugars in HC1 and HC2 was analyzed by the phenol-sulfuric acid method expressed as glucose equivalents (Dubois *et al*., 1956). Briefly, 200 µL sample was reacted for 15 min with 10 µL phenol (80%) and 1 mL concentrated sulfuric acid. The absorbance at 490 nm was measured after 15 min of hot water bath at 100 °C.

### 2.6. Root ruthenium red staining

The ruthenium red staining were prepared by the method of Ballance *et al*., (2012), respectively (detail can be found in the supplementary data).

### 2.7. Aluminum extraction and quantitation in CW materials

30 mg of CW and cellulose were extracted in 1.2 mL of 2 M HCl for 24 h with occasional shaking. Similarly, Al content in each CW component extract (WSP, ASP, CSP, HC1, and HC2) was extracted by 2 M HCl (1:1, v/v) for 24 h as described above. The Al content in each liquid extract was diluted as necessary and quantified by ICP-OES.

### 2.8. Transmission electron microscope (TEM) analysis

The root tips (0-5 mm) of seedlings were cut and quickly placed in 2.5% glutaraldehyde solution at 4 °C for 12 h, and then washed four times (15 min each time) with 0.1 M phosphate buffer (pH=7.4). Then, the tissue blocks were fixed in 1% osmium tetroxide (OsO_4_) for 2-3 h and rinsed three times with 0.1 M PBS (pH 7.4) for 45 min, and then dehydrated three times with 50%, 70%, 90% ethanol and once with the mixture of 90% ethanol and 90% acetone (1:1, v/v) for 15 min, respectively. Finally, the embedded samples were washed with 100% acetone. The ultrathin sections were stained with 2% uranyl acetate and lead citrate solution, and then observed and photographed under JEM-100CXIITEM (Kong *et al*., 2013).

### 2.9. Measurement of activity of pectin methylesterase (PME) and degree of methylation (DM) of pectin

The activities of PME in roots were estimated by the PME kit followed by the instructions of the manufacturer (Shanghai You Xuan Biotechnology Company Limited). The DM of pectin was measured by the Louvet *et al*., (2011) method and absorbance was recorded colourimetry at 620 nm. The methanol resulting from saponification and formaldehyde was calculated by Anthon & Barrett (2004) and Yang *et al*., (2008).

### 2.10. Determination of H_2_O_2_ and MDA contents, total antioxidant capacity (T-AOC), and plasma membrane (PM) H^+^-ATPase activity

The MDA content was carried out in root based on the method of Vos *et al*., (1991) with slight modifications. The H_2_O_2_ content, total antioxidant capacity (T-AOC), and PM H^+^-ATPase activity in root were measured by commercially available ELISA H_2_O_2_ kits, T-AOC kits, and PM H^+^-ATPase kits, respectively, and followed the guidelines of manufactures (Shanghai You Xuan Biotechnology Company Limited).

### 2.11. Fourier-transform infrared spectroscopy (FTIR) analysis of CW

The CW discs for FTIR analysis were prepared with the homogeneous mixture (KBr at the ratio of root CW, 1:100, m/m) by Graseby-Specac Press. Infrared spectra of CWs in the range of 4000-400 cm^-1^ (64 scans/samples, resolution 4 cm^-1^) were analyzed by spectrophotometer (Vertex 70, Brooklyn Instrument, USA).

### 2.12. X-ray photoelectron spectroscopy (XPS) and scanning electron microscope-energy dispersive x-ray spectrometer (SEM-EDS) analysis of Al content in CW surface

The CW sample (through a 0.150 mm sieve) was measured by XPS. The tests were carried out under vacuum conditions with a pressure of 3×10-10 mbar. The survey scan had a resolution of 1 eV. The high-resolution spectra were recorded with a pass energy of 30 eV and an energy step size of 0.1 eV for the scan of Al2p.

The microstructure of at the sample was analyzed using a Zeiss GeminiSEM Family Sigma 500 FEG-SEM (Field Emission Gun-Scanning Electron Microscope) equipped with an EDS (Energy Dispersive x-ray Spectroscopy) analyzer.

### 2.13. Transmission electron microscope-energy dispersive x-ray spectrometer (TEM-EDS) analysis of Al content in intercellular space, cytoplasm, and vacuoles

The procedure of the root tip (0-5 mm) section was similar to that of TEM. Low-dose TEM imaging was performed on a JEOL JEM2100 microscope operated at 200 kV (Cs 1.0 mm, point resolution 0.23 nm). Samples for TEM were dispersed in acetone. A droplet of the suspension was transferred onto a carbon-coated copper grid. Images were recorded with a Gatan Orius 833 CCD camera (resolution 2,048 × 2,048 pixels,pixel size 7.4 μm) under low-dose conditions 41.

### 2.14. Quantitative real-time PCR (qRT-PCR)

Total RNA in the root tips (0-10 mm) was extracted using Eastep^®^ Super Total RNA Extraction Kit (Shanghai Promega Biological Products Ltd). Gene-specific primers were designed by the National Center for Biotechnology Information (NCBI) and are shown in Table S2. First-strand cDNA was synthesized by ReverTra Ace^®^ qPCR RT Master Mix with gDNA Remover Kit (Toyobo Co., LTD. Life Science Department OSAKA Japan). Reactions were carried out by Hieff^®^ qPCR SYBR^®^ Green Master Mix Kit (Shanghai Yeasen Biotech Co., Ltd) and the manufacture’s guidelines followed procedures in a 384-well real-time PCR Detection mixtures consisted of 5 µL of master mix, 0.2 µL forward primer (10 µM), 0.2 µL reverse primer (10 µM), 2 µL of cDNA template, and sterile ultra-pure water was added to a final volume of 10 µL. The optimal PCR amplification procedure indues: Pre-denaturation at 95 °C for 5 min, denaturation at 95 °C for 10 s, annealing and extension for 20 s at 60 °C and 72 °C, respectively. There were three replicates in each treatment. The expression levels of related genes were calculated by 2^-ΔΔCT^ method (Livak & Schmittgen, 2001).

### 2.15. Data Statistics and Analysis

All statistical analysis were carried out by statistical software (SPSS 16.0, Chicago, IL, USA). The data were examined with one-way analysis of variance (ANOVA) followed by Duncan’s tests. The FTIR data were subjected to OMNIC 8.2 (Thermo Fisher Scientific Inc., USA) and Origin 8.6 software (Origin Lab Corporation, USA), respectively.

## 3. Results

### 3.1. Effects of B on growth parameters of seedlings under Al stress

In the absence of Al, the seedlings treated with B at 5 (B5), 10 (B10), and 50 µM (B50) significantly increased the plant growth compared to 0.1 µM B (B0.1). –Al B50 treatment did not increase the plant growth-related parameters compared with –Al B10 (Fig. 1). Aluminum toxicity had a detrimental effect on plant growth characteristics, the plant height, root length, and shoot DW were significantly decreased by +Al B0.1 treatment compared to –Al B0.1 (Fig. 1c-h). Furthermore, the chlorosis was observed at the top new leaves, which manifested starting at the tips along the margins to form scorched leaves. Besides, the root tips and lateral roots showed obvious enlargement symptoms (Fig. 1b). While in the presence of Al, increased B levels in nutrient solutions significantly increased the plant growth, especially root length (Fig. 1c-h). Similar results are observed in Fig. S1 that application of 10 and 50 µM B can also alleviate the inhibitory effect of Al on seedlings biomass.

**Fig. 1.**
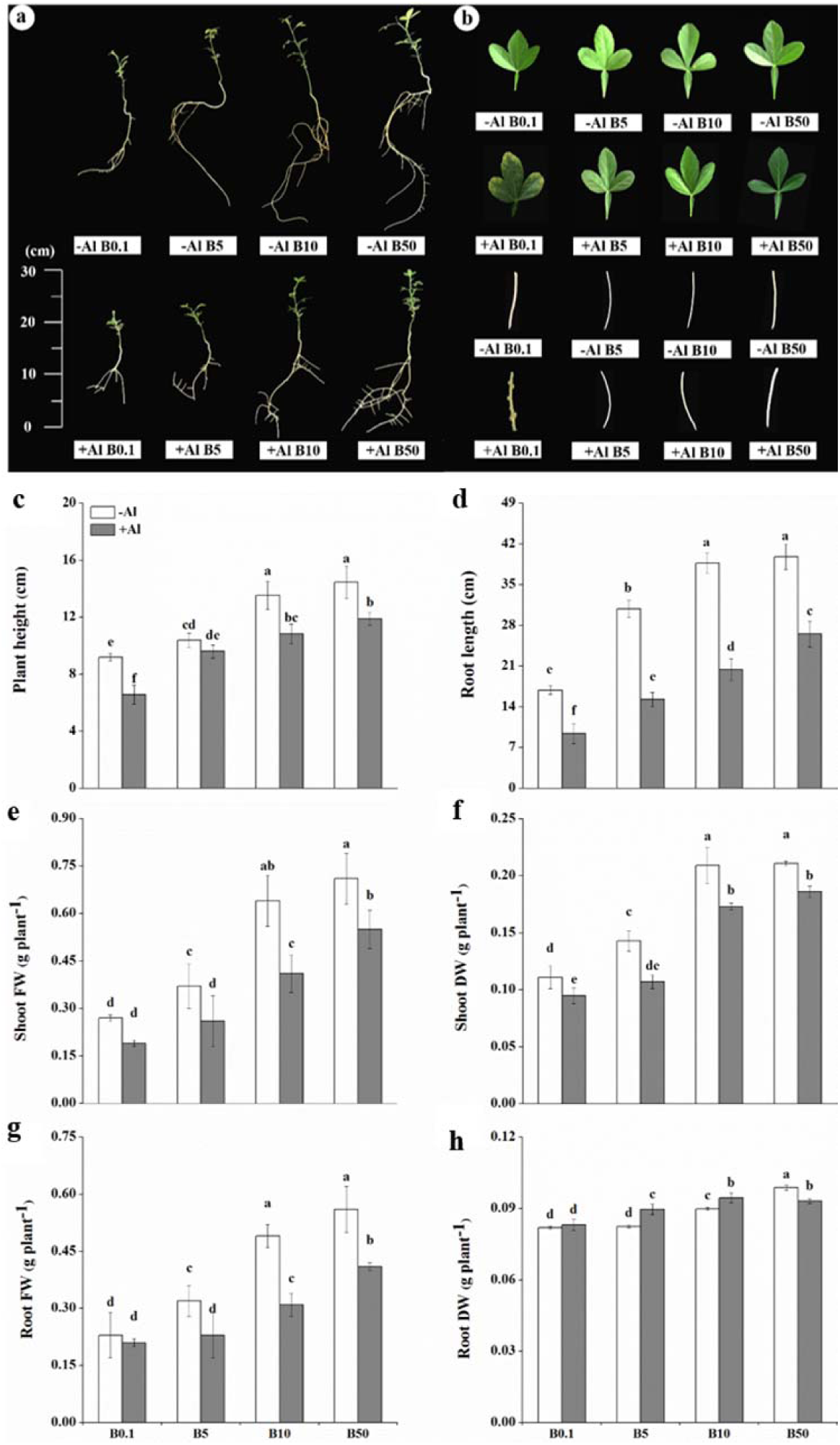
Effects of different B concentrations on the growth (a), symptoms (b), plant height (c), root length (d), shoot (stem + leaf) fresh weight (FW) (e), shoot dry weight (DW) (f), root FW (g), and root DW (h) of trifoliate orange rootstock under Al stress. The experimental treatments consisted of four B levels (0, 5, 10, and 50 µM as H_3_BO_3_), and two Al levels (0 and 300 µM). Fig-b: The leaves and roots represent the new apical leaves and root tips (0-10 mm), respectively. Bars are means of three replicates ± SD. Different letters (a, b, c, d) in each sub-figure represent significant differences at (*P* < 0.05).

### 3.2. Effects of B on B and Al content in the root, Al content in root tip under Al stress

The B concentration in roots was obviously increased after treatment with different B levels, regardless of Al treatment (Fig. 2a; Table S1). Aluminum content in root and root tips was remarkably increased under B0.1 treatment. Under Al stress, the Al content in root and root tips decreased gradually with the increase of B concentration (Fig. 2b-c). To further verify the practical application of B to reduce Al content in plants, we conducted a similar hydroponics experiment. The result in Table S1 showed that B obviously reduced Al content in citrus roots and root tips compared with the Al treatments with B0.1.

**Fig. 2.**
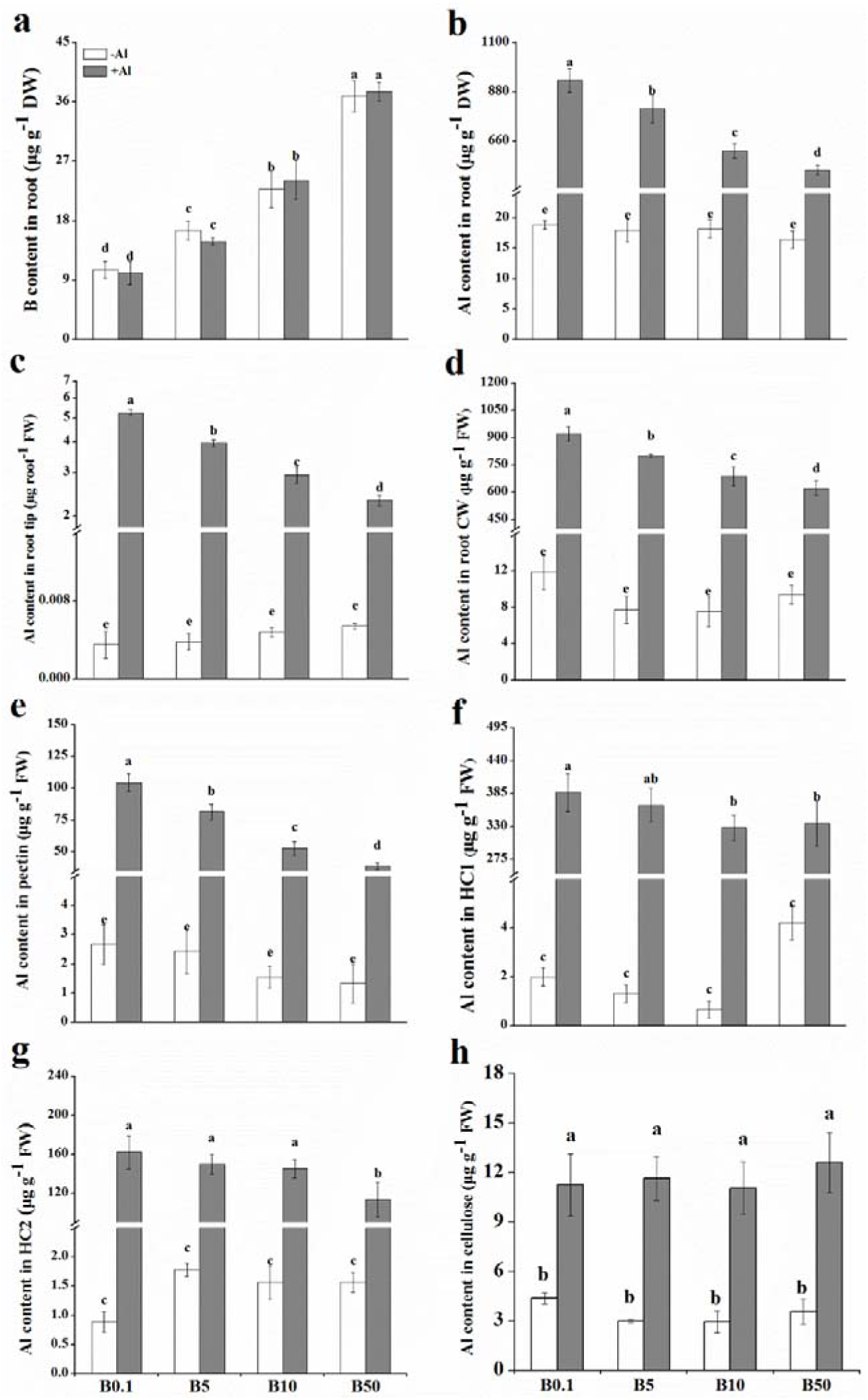
Effects of different B concentrations on the B content in the root (a), Al content in the root (b), Al content in the root tip (c), Al content in root CW (d), Al content in pectin (e), Al content in HC1 (f), Al content in HC2 (g), and Al content in cellulose (h) of trifoliate orange rootstock under Al stress. Bars are means of three replicates ± SD. Different letters (a, b, c, d) in each sub-figure represent significant differences at (*P* < 0.05).

### 3.3. Effects of B on Al content in root CW under Al stress

Aluminum accumulation in CW accounted for the most part of total Al in root: 79.60-87.33% of Al content in root CW was present (Fig. 2d), the addition of B reduced Al accumulation in root CW (Fig. 2d; Table S1). XPS was used to analyze the characteristics of Al on root CW under different treatments. The result in Fig. 3 showed that Al existed mainly in Al_2p_ and Al_2p3/2_ forms. Al_2p_ is Al/(-CH_2_CH(C(O)OH)-), which binding energy is 73.9eV (Dekoven & Hangans, 1986). Al_2p3/2_ is Al_2_O_3_ and Al(OH)_3_ with the binding energy of 74.9 and 75.9eV, respectively (Mcguire *et al*., 1973). There was no significant effect on the atomic ratio of Al between different B treatments without Al (Fig. 3a-c; Table S3). +Al B0.1-treated root CW exhibited a higher Al atomic ratio level (from 0.56 to 1.77) in comparison to –Al B0.1. In the presence of Al, Al atomic ratio in root CW declined by 33.90 and 37.85% under B10 and B50 treatment, respectively compared to B0.1 (Fig. 3d-f; Table S3). Meanwhile, SEM-EDS data visually reflected the accumulation of Al in the CW under different treatment conditions. Point scan data showed that the Al absorption edge energy intensity of the treated groups without Al was similar and approached zero (Fig. S2a-c). Aluminum-treated roots dramatically increased the energy intensity of Al absorption edge, along with the increase of the weight ratio and atomic ratio of Al element, especially under B0.1 treatment (Fig. S2a, d). Al absorption edge energy intensity, weight ratio and the atomic ratio of Al in root were observed to be lower with 10 and 50 µM B supplementation than with 0.1 µM B during Al exposure (Fig. S2d-f). Various results demonstrated that applied B exerted beneficial effects on the reduction of Al accumulation in root CW as well.

**Fig. 3.**
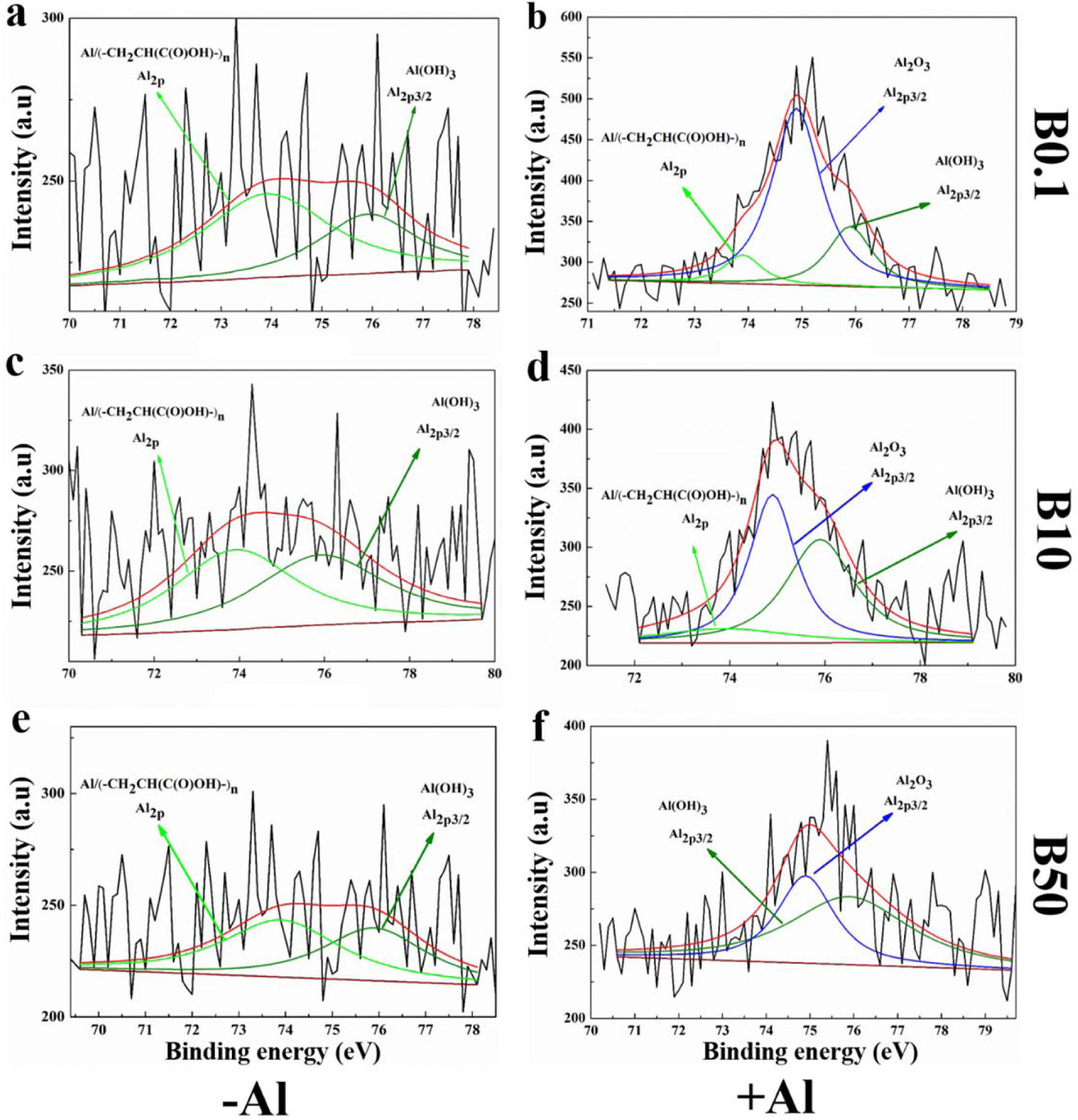
Effects of B on the deconvoluted high-resolution XPS Al spectrum of CW in trifoliate orange rootstock root under Al stress. Fig-a: Plants treated with 0 µM Al and 0.1 µM B (–Al B0.1); Fig-b: Plants treated with 0 µM Al and 10 µM B (–Al B10); Fig-c: Plants treated with 0 µM Al and 50 µM B (–Al B50); Fig-d: Plants treated with 300 µM Al and 0.1 µM B (+Al B0.1); Fig-e: Plants treated with 300 µM Al and 10 µM B (+Al B10); Fig-f: Plants treated with 300 µM Al and 50 µM B (+Al B50).

### 3.4. Effects of B on Al content in root CW components under Al stress

Aluminum contents in different components of CW had no significant effect in the absence of Al (Fig. 2e-h). Indeed, the Al content in HC1 fraction of CW was higher than that in HC2 under Al stress, both of them were higher than that in pectin fraction, and the proportion of Al content in cellulose fraction of CW was the smallest (Fig. 2e-h). The seedlings exhibited enhanced Al contents in B0.1-treated root CW pectin after Al exposure when compared to Al-untreated treatment (Fig. 2e). However, under Al stress, levels of root pectin Al content were 22.08%, 49.47%, and 62.85% under B5, B10, and B50 treatments lower than B0.1, respectively (Fig. 2e).

We further investigated the content of Al in different forms of pectin, the distribution of Al content in pectin was ASP > WSP > CSP (Table S1). Aluminum contents in three kinds of pectin were increased significantly under Al stress irrespective of B treatments and the effects were more pronounced in B0.1-treated roots. The addition of 10 and 50 µM B resulted in lower Al content than at 0.1 µM B in the presence of Al, notably, ASP had the most obvious decrease in Al content. +Al B50 did not cause remarkable effects on Al uptake in CSP compared to +Al B10 (Table S1).

Moreover, Al contents in HC1, HC2, and cellulose fraction were increased in +Al B0.1 treatment compared to –Al B0.1 (Fig. 2f-h). In the presence of Al, B5 did not affect Al content in HC1 compared to B0.1, whereas it decreased by 15.38% (from 387.11 to 327.56 µg g^-1^) in B10 and 13.32% (from 387.11 to 335.56 µg g^-1^) in B50 treatment (Fig. 2f). Interestingly, Al contents in both HC2 and cellulose fractions were unaffected by exposure to different levels of B compared to B0.1 (Fig. 2g-h), except the HC2, Al contents were reduced in +Al B50 compared to +Al B0.1 (Fig. 2g).

### 3.5. Effects of B on CW components content under Al stress

Aluminum-treated roots significantly increased the pectin content (indicated by the uronic acid content in pectin), HC1 and HC2 contents (indicated by the total sugar content in HC1 and HC2) and increased by 1.64, 1.54, and 1.50-fold under B0.1 treatment, respectively (Fig. 4g-k). However, B5, B10, and B50 applications decreased Al-induced increase of pectin content (Fig. 4g). Ruthenium red dyeing characterizes the content of pectin in root CW, +Al B0.1 treatment had pink color in root tips (Fig. 4d), whereas, the treatment of +Al B10 and +Al B50 resulted in the reduced content of pectin in the roots with a faded pink staining (Fig. 4-f), further authenticating that B considerably decreased the pectin content in root CW. In addition, the content of HC1 was unaffected with B addition at 0.1, 5, and 10 µM under exposure to Al, while HC1 content was decreased significantly only under +Al B50 treatment (Fig. 4j). Moreover, in the presence of Al, B addition (B5, B10, and B50) reduced the HC2 content compared to B0.1. Interestingly, there was no significant effect on the content of HC2 among B5, B10, and B50 treatments under Al stress (Fig. 4k).

**Fig. 4.**
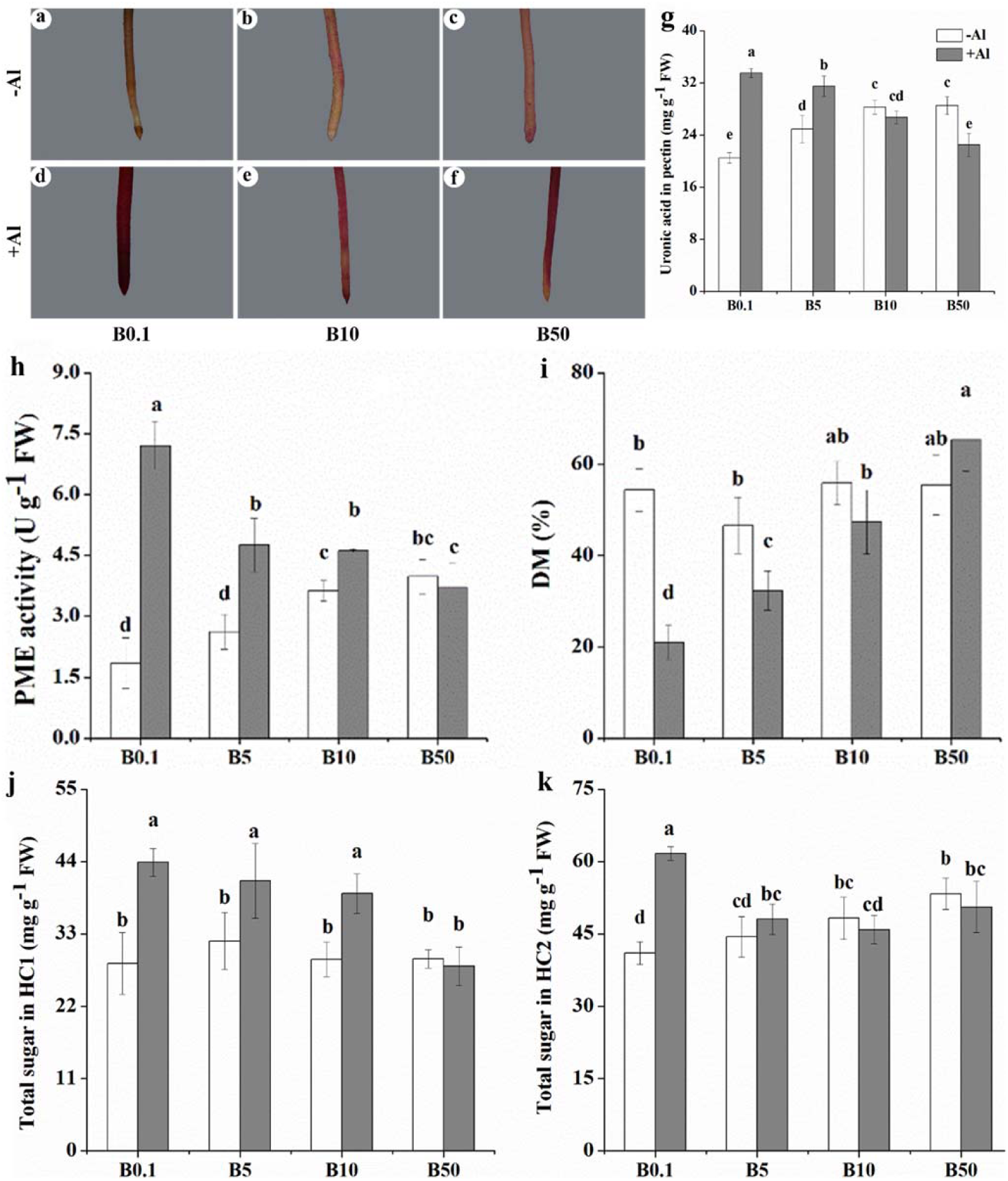
Effects of different B concentrations on ruthenium red staining of the root tips (a-f), uronic acid content in pectin (g), PME activity (h), degree of methylation (DM) (i), total sugar in HC1 (j), and total sugar in HC2 (k) in the root of trifoliate orange rootstock under Al toxicity. Fig-(a-f): The distribution of pectin indicated by ruthenium red (red fluorescence) staining. Bars are means of three replicates ± SD. Different letters (a, b, c, d) in each sub-figure represent significant differences at (*P* < 0.05).

The contents of the three types of pectins are shown in Fig. 5. The content of ASP was more abundant than that of WSP, and CSP was the least in root CW. There was no significant effect of different B concentrations on the contents of three kinds of pectins without Al treatment (Fig. 5a). Supplementation of Al increased the contents of WSP and ASP while decreased the CSP contents in root CW under B-deficiency. The WSP and ASP contents in root CW were found to be lower under 10 and 50 µM B supplementation compared to 0.1 µM B during Al exposure, and the decline was more marked in ASP. It was observed that there was no significant difference in CSP content in the treatment of +Al B0.1, +Al B10, and +Al B50 (Fig. 5a).

**Fig. 5.**
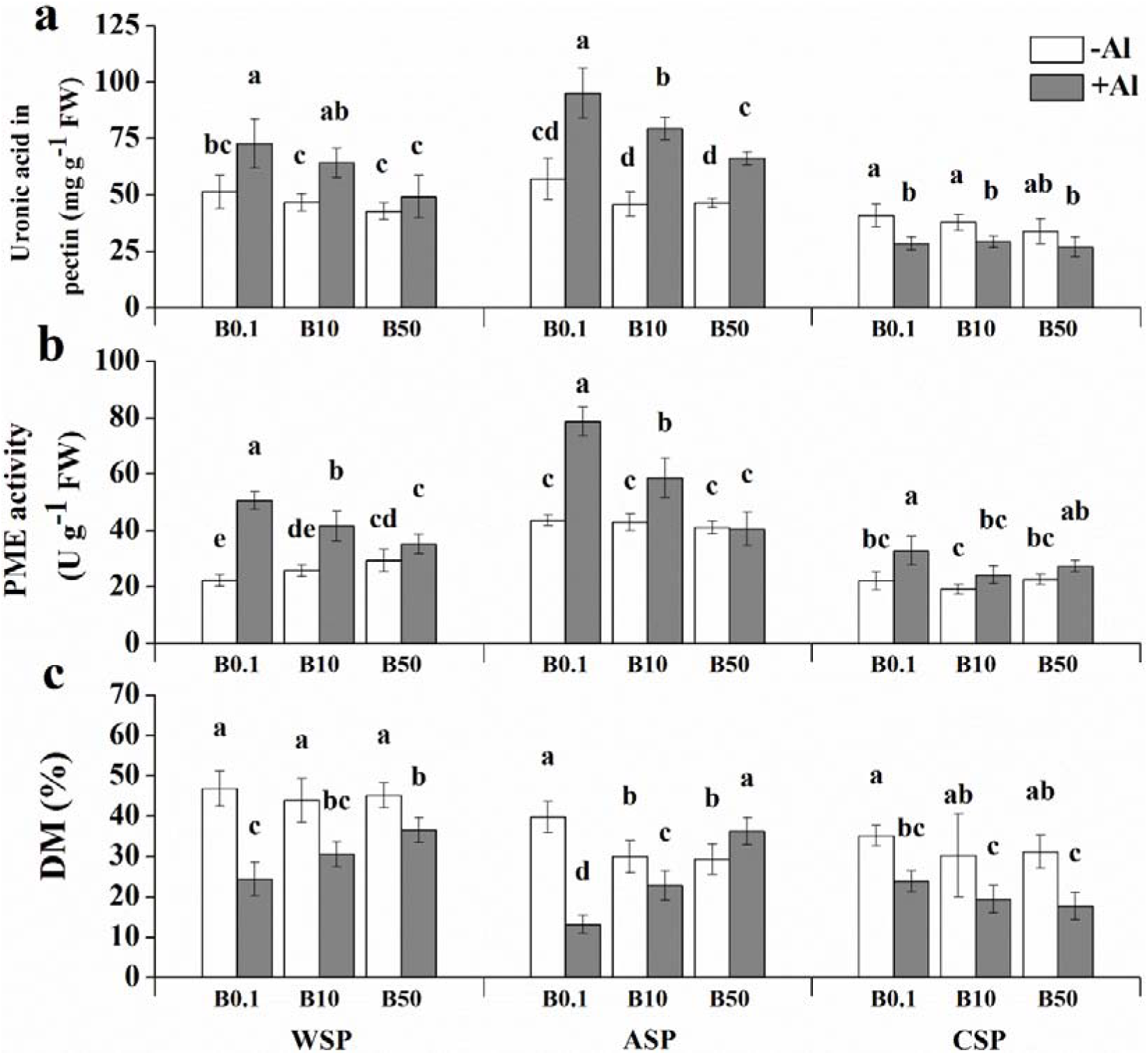
Effects of B on uronic acid content (a), PME activity (b), degree of methylation (DM) (c) in different pectin forms of trifoliate orange rootstock root under Al toxicity. WSP: water-soluble pectin; ASP: alkali-soluble pectin; CSP: chelator-soluble pectin. Bars are means of three replicates ± SD. Different letters (a, b, c, d) in each sub-figure represent significant differences at (*P* < 0.05).

### 3.6. Effects of B on pectin properties under Al stress

Aluminum toxicity increased PME activity while decreased the DM in root under B0.1 treatment (Fig. 4h-i). PME activity in +Al B0.1-treated root was about 3.90-fold with respect to –Al B0.1 treatment (Fig. 4h). Nevertheless, +Al B5, +Al B10, and +Al B50 treatments significantly decreased PME activity in the root; the DM showed the opposite trend, i.e. obviously increased compared to +Al B0.1 (Fig. 4i). Similarly, the PME activity in WSP, ASP, and CSP amplified under Al-treated roots regardless of B concentration and particularly under B-deficiency (Fig. 5b). Compared to 0.1 µM B, 10 and 50 µM B substantially dropped the PME activity in roots under Al toxicity. PME activity in WSP decreased by 17.79 and 30.39% at 10 and 50 µM B treatments, respectively, and declined by 25.47 and 48.55% in ASP at 10 and 50 µM B treatments under Al stress (Fig. 5b). The DM of WSP, ASP, and CSP drastically inhibited after Al employing with B-deficiency. In the presence of Al, the DM of WSP and ASP were intensified under B10 and B50 treatment in comparison to B0.1. In contrast, the DM of CSP had a slight downward trend with the increase of B concentration, although the difference was not significant (Fig. 5c).

### 3.7. FTIR analysis of root CW

Effects of Al on changes of CW functional structure and chemical components of CW involving in the binding of Al were characterized by FTIR analyses in the range of 4000-400 cm^-1^ (Fig. S4). The spectral results instructed that the positions of absorption peaks did not create, while the intensities of absorption peaks were greatly influenced by Al toxicity. The peak values at 2928 and 2857 cm^-1^ representing the saturated C-H stretching related to protein, cellulose, and pectin were enhanced in the treatment of +Al B0.1 compared to –Al B0.1 treatment. The intensities of absorbance located at 1731 cm^-1^ corresponding to C=O stretching from -COOR mainly pectin in Al toxicity were more evident than control. Root CW spectra indicated higher intensities of absorbance at 1379 and 1248 cm^-1^ in the +Al B0.1 treatment, reflecting the saturated C-H deformation in cellulose. The wavelengths situated at 1025 cm^-1^ demonstrating the characteristics of the polysaccharide region -CH deformation or C-C, C-O stretching, were enhanced in Al-treated root CW. Whereas B application at 10 and 50 µM concentrations showed a significant decline in 1731, 1379, and 1248 peaks values related to pectin and cellulose compared to +Al B0.1 (Fig. S4).

### 3.8. Transmission electron microscope (TEM) of root tips

The ultrastructure of roots and the changes in the subcellular level were observed by TEM (Fig. S3). In the absence of Al, the TEM micrographs showed that compared with B0.1 treatment, cells under B5 and B10 treatment were closely arranged and showed regular shape, especially in B10 treatment (Fig. S3a-c). The cells were flattened under –Al B50 treatment (Fig. S3d). +Al B0.1 treatment resulted in irregular cell characteristics, besides, CW was thickened and starch granule depositions were obvious (Fig. S3e, m). Boron (B5, B10, and B50) application reduced the CW thickness (Fig. S3q), as well as a reduced the accumulation of starch grains, which showed the clear, and well-arranged structure (Fig. S3f-h, n-p) indicating that B had a significant mitigation effect on Al toxicity.

### 3.9. Effects of B on Al content in intercellular space, cytoplasm, and vacuoles under Al stress

The Al content in the intercellular space (zone 1), cytoplasm (zone 2) and vacuole (zone 3) of root tips (0-5 mm) were further determined by TEM-EDS (Fig. 6a-c). In the presence of Al, the B content in different parts (indicated by the weight ratio) increased correspondingly with the increase of B concentration. Interestingly, the Al content in these three parts (indicated by the weight ratio) gradually decreased in +Al B10 and +Al B50 treatment compared to +Al B0.1 (Fig. 6d-l). Therefore, through further analysis of the proportion of Al content in intercellular space, cytoplasm, and vacuole to total Al content in root tips (Fig. S3), B10 and B50 treatment increased the proportion of Al content in intercellular space and vacuole in comparison to B0.1 under Al stress, whereas, the lower proportion of Al content in cytoplasm was observed in B10 and B50 treatment (Fig. S3).

**Fig. 6.**
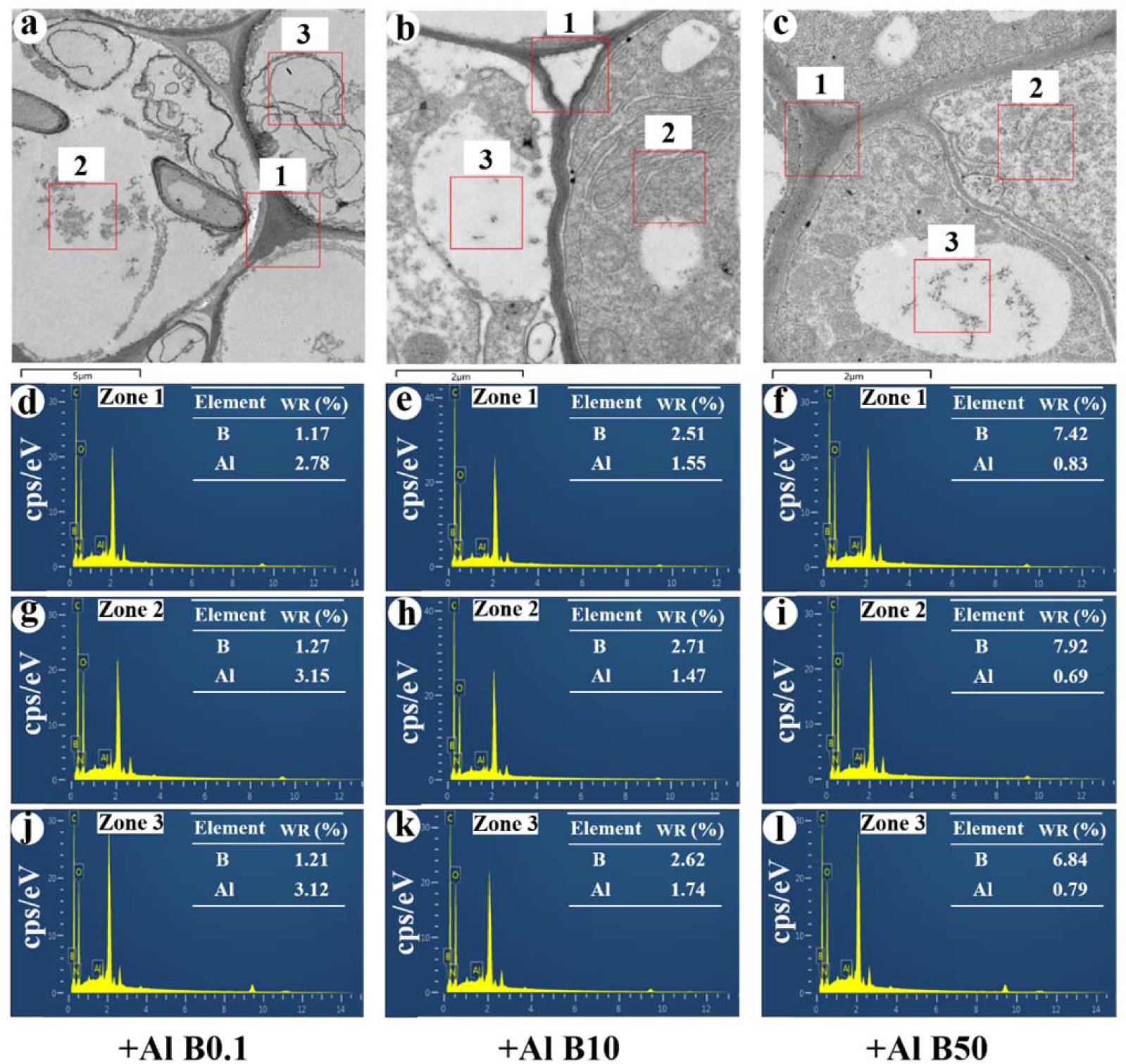
Effects of B on Al content in intercellular space, cytoplasm, and vacuoles of the trifoliate orange rootstock root tips under Al toxicity. Plants treated with 300 µM Al and 0.1 µM B (+Al B0.1); Plants treated with 300 µM Al and 10 µM B (+Al B10); Plants treated with 300 µM Al and 50 µM B (+Al B50); WR: weight ratio.

### 3.10. Effects of B on Al uptake and translocation-related gene expression under Al stress

Aluminum treatment obviously inhibited the expression of *ALS1* while induced the *STAR1* expression compared to the control, although *PME* showed an upward trend and *NRAT1* showed a downward trend in +Al B0.1 treatment in comparison to – Al B0.1, there was no significant difference (Fig. 7). In the presence of Al, B10 and B50 application further reduced the *PME* and *NRAT1* expression compared to B0.1 treatment, while +Al B10 and +Al B50 up-regulated the *ALS1* and *STAR1* expression (Fig. 7a-d).

**Fig. 7.**
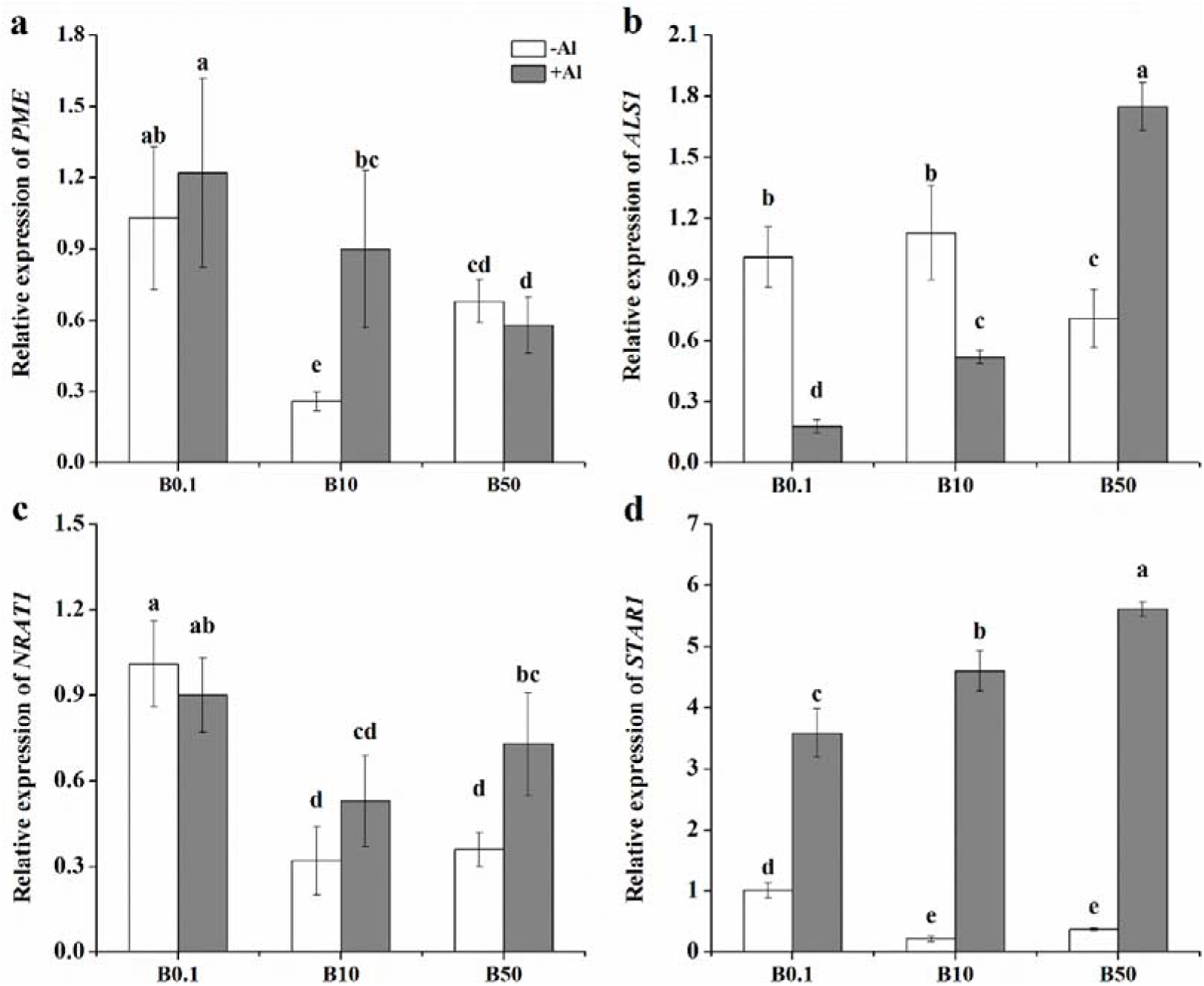
Effects of B on the expression of *PME* (a), *ALS1* (b), *NRAT1* (c), and *STAR1* in the trifoliate orange rootstock root tips under Al toxicity. Bars are means of three replicates ± SD. Different letters (a, b, c, d, e) in each sub-figure represent significant differences at (*P* < 0.05).

### 3.11. Effects of B on H_2_O_2_ and MDA contents, T-AOC, and PM H^+^-ATPase activity in roots under Al stress

The H_2_O_2_, MDA contents, and T-AOC of the root under the +Al B0.1 treatment were higher than that of the –Al B0.1, while PM H^+^-ATPase activity in the Al-treated root was lower than that of the without Al treatment under B-deficiency (Fig. S6). The H_2_O_2_, MDA contents, and T-AOC in roots were substantially decreased both at 10 and 50 µM B with Al in comparison to 0.1 µM B (Fig. S6a-c). 10 and 50 µM B substantially enhanced the PM H^+^-ATPase activity under Al toxicity and improved by 1.37 and 1.71-fold, respectively (Fig. S6d).

## 4. Discussion

Boron is indispensable for plant growth and in the mitigation of Al toxicity (Zhu *et al*., 2019). In the previous study, we have explored mechanisms by which B could regulate Al stress in the rape seedlings (Yan *et al.*, 2018b) and citrus (Yan *et al*., 2018a; Riaz *et al*., 2018). When seedlings were exposed to Al stress, plant growth-related parameters such as plant height, root length, and root elongation were inhibited (Fig. 1; Fig. S1a-d). Moreover, Al toxicity symptoms such as enlargement of root tips and yellowing of upper leaf tips were observed (Fig. 1b). The CW of root tips was also thickened and more starch grains were observed by TEM (Fig. S3e, m). Meanwhile, the intensities of 2928, 2857, 1379 and 1248 cm^-1^ absorption peaks related to cellulose were increased significantly under Al stress (Fig. S4). This indicates that the accumulated cellulose inhibits CW structure, viscosity and elasticity, thus inhibits root elongation (Ma *et al.*, 2004; Zhou *et al.*, 2014). We found that high B (B10 and B50) decreased cellulose content at absorption peaks intensities of 2928, 2857, 1379, and 1248 cm^-1^ (Fig. S4), thereby reduced CW thickness (Fig. S3n-p), meanwhile, B application significantly decreased the H_2_O_2_ and MDA content and T-AOC under Al stress (Fig. S6a-c), illustrating B reducing the Al-induced accumulation of ROS and abating the damage to plants by regulating root oxidation ability, thereby promoting root elongation (Fig. 1; Fig. S1). The B contents in roots were supplemented with the increase of B levels under Al stress, interestingly, Al treatment had no significant effect on B uptake and accumulation in roots (Fig. 2a; Table S1), which is in agreement with our previous research (Yan *et al.*, 2018c) indicating that (1) a non-obviously effect of Al on B adsorption in roots; (2) the accumulation of Al in roots was the main driving force to inhibit root growth and (3) B has a vital protective role in reducing Al-induced inhibition to plants.

The amount of Al accumulation in roots was positively correlated with Al toxicity (Wang *et al.*, 2017). In the present study, Al significantly enhanced Al content in root and root CW (Fig. 2b-d; Table S1), which are in agreement with the findings of Chang *et al.*, (1999) that more than 80% Al binds to root CW (Fig. 2d). Boron supply decreased Al content in the root (Fig. 2b), more specifically reduced Al content in root CW (Fig. 2b-d; Table S1). Simultaneously, XPS in conjunction with SEM-EDS further proved that B application decreased the atomic ratio of Al in root CW (Fig. 3d-f; Fig. S2d-f), which suggested that the alleviating effect of B on Al toxicity can be due to reduced accumulation of Al in root CW. Although absorption and accumulation of Al in CW have been intensively studied, their roles in CW components remain to be unclear. In this study, we found that most of the Al accumulated in HC1 (Fig. 2f) and a similar study also found in alfalfa (Wang *et al.*, 2017) that a large amount of Al was accumulated in HC1 and significantly higher than that in pectin, showing that Al^3+^ interacted with hemicellulose polysaccharides and most of the Al accumulated in HC1 (Yang *et al.*, 2011a). Similar conclusions were drawn in *Arabidopsis* research that the increase of Al binding in HC1 component inhibited root elongation (Zhu *et al.*, 2012). Meanwhile, Al^3+^ binding in pectin can also inhibit plant growth (Li et al. 2017; Yan *et al.*, 2018a), interestingly, different B levels supply decreased Al content in pectin gradiently (Fig. 2e), especially in WSP and ASP (Table S1), but was not statistically significant in Al content in HC1 and HC2, except for +Al B50 treatment (Fig. 2f-g), suggesting that B addition on the ability of pectin binding Al is more obvious than other components, and more specific action in ASP, which in line with the conclusion that ASP is the predominant target of Al accumulation in CW (Li *et al.*, 2017). A large amount of pectin, HC1, and HC2 were observed under +Al B0.1 treatment (Fig. 4g-k) indicating pectin and hemicellulose are different in the aspect of the binding capacity of Al, higher Al adsorption in CW was positively correlated with its higher HC1 and HC2 contents (Wang *et al.*, 2017). Alike, the addition of 10 and 50 µM B reduced pectin content (Fig. 4g), the root tips (0-10 mm) were further stained with ruthenium red, demonstrating that the pectin levels (stained reddish) were decreased by B under Al stress conditions (Fig. 4d-f). In addition to pectin content, the DM of pectin also affects the binding capacity and sensitivity of Al in many plants (Horst *et al.*, 2010), which is controlled by PME, and the Al binding capacity of CW is negatively responsible for the methylation (Eticha *et al.*, 2005). In our study, we further analyzed the pectin properties and found that with the increase of B levels, the content of ASP and PME activities in ASP severely decreased while DM of ASP apparently increased gradiently (Fig. 5a-c), meanwhile, the *PME* expression associated with PME activity were down-regulated by B application under Al stress conditions (Fig. 7a), indicating that B crosslinking with the ASP and promoted Al immobilization in pectins with a higher DM by reducing PME activity, and then decreased the Al enrichment in CW (Zhu *et al.*, 2018). More importantly, B reduced the absorption peak strength at 1731 cm^-1^ (characteristic of the C=O stretching vibration from -COOR in pectin) (Fig. S4), which consequently decreased the carboxyl groups in pectin, thus excluded CW bound Al (Yan *et al.*, 2018a). In addition, the role of B in promoting the demethylation of pectin is related to the surface pH (Li *et al.*, 2017), PME activity in roots was positively correlated with apoplast pH and intracellular H^+^ level (Denès *et al.*, 2000; Jolie *et al.*, 2009). The PM H^+^-ATPase is a proton pump, and its hydrolysis is accompanied by transmembrane H^+^ transport (Volkmar & Hu, 1998). Aluminum stress reduced the PM H^+^-ATPase activity (Fig. S6d), which induced lesser H^+^ transfer from internal cells to apoplastic space and a reduction of H^+^ secretion from root tips (Wang *et al.*, 2017). The increase of PM H^+^-ATPase activity by B (Fig. S6d) can promote the excretion of H^+^ from intracellular and reduce the apoplast pH, thus weakening the Al-induced intracellular acidic environment and stabilizing the cell structure (Ahn *et al.*, 2001). The decrease of intracellular H^+^ content by B inhibited PME activity in turn (Fig. 4h). These results suggested that the effect of B on Al-toxicity might be implemented by maintaining H^+^ balance between external and internal cells.

Plants often regulate Al^3+^ transport in cells to reduce Al toxicity. NRAMP ALUMINIUM TRANSPORTER1 (*NRAT1*) transports Al^3+^ from the outside of cells to inside of cells, and increase the concentration of Al in the cytoplasm and inhibits root growth (Zhu *et al*., 2018). +Al B10-treated roots significantly decreased the *NRAT1* expression (Fig. 7c) and increased the proportion of Al in intercellular space (Fig. S5) compared to +Al B0.1 treatment, illustrating that B reduced the amount of Al into the cells. Simultaneously, Huang *et al*., (2012) reported that *OsALS1* encodes a half-size ABC transporter, involved in quarantine rice cytoplasmic Al into the vacuole. B supply (50 µM) enhanced the Al-induced decreases in the expression of *ALS1* (Fig. 7b), accompanied by lesser cytoplasmic Al ration and higher vacuole Al ratio of root tips after B10 and B50 addition (Fig. S5), and yielded similar results in rice that B application could increase the *ALS1* expression (Zhu *et al*., 2019), suggesting that B could provoke the transfer of Al from the cytoplasm to the vacuoles, which showing low Al toxicity to seedlings. In addition, the expression of *OsSTAR1* and *OsSTAR2*, plays an important role in rice Al tolerance (Zhu *et al*., 2018; Zhu *et al*., 2019), and functions as a transporter of the ATP binding cassette, witch transports UDP-glucose from the cytoplasm to CW, thus reducing the deposition of Al in CW (Huang *et al*., 2009). In the current study, *STAR1* expression increased in response to Al stress, and treatment with B10 and B50 further induced the expression of *STAR1* under Al stress (Fig. 7d), which was accompanied by a decrease in Al content in CW (Fig. 2d), further authenticating that B alleviated the Al toxicity of citrus by reducing Al content in CW.

## 5. Conclusions

As summarized in the proposed model above (Fig. 8), the accumulation of Al in CW plays a major role in Al toxicity of citrus, and the most of Al in CW chiefly binds to HC1 fraction. The protection of B supply under Al stress is mediated by a mechanism that enhanced the PM H^+^-ATPase activity, thus promoting H^+^ efflux and decreased apoplast pH, meanwhile, B application decreased pectin synthesis and PME activity, thereby decreasing Al binding site (carboxyl group) in pectin and reduced Al fixation of the CW that ultimately alleviated Al toxicity. Moreover, B functions by suppressing the absorption of Al in intracellular and accelerating the compartmentalization of Al from the cytoplasm into the vacuole.

**Fig. 8.**
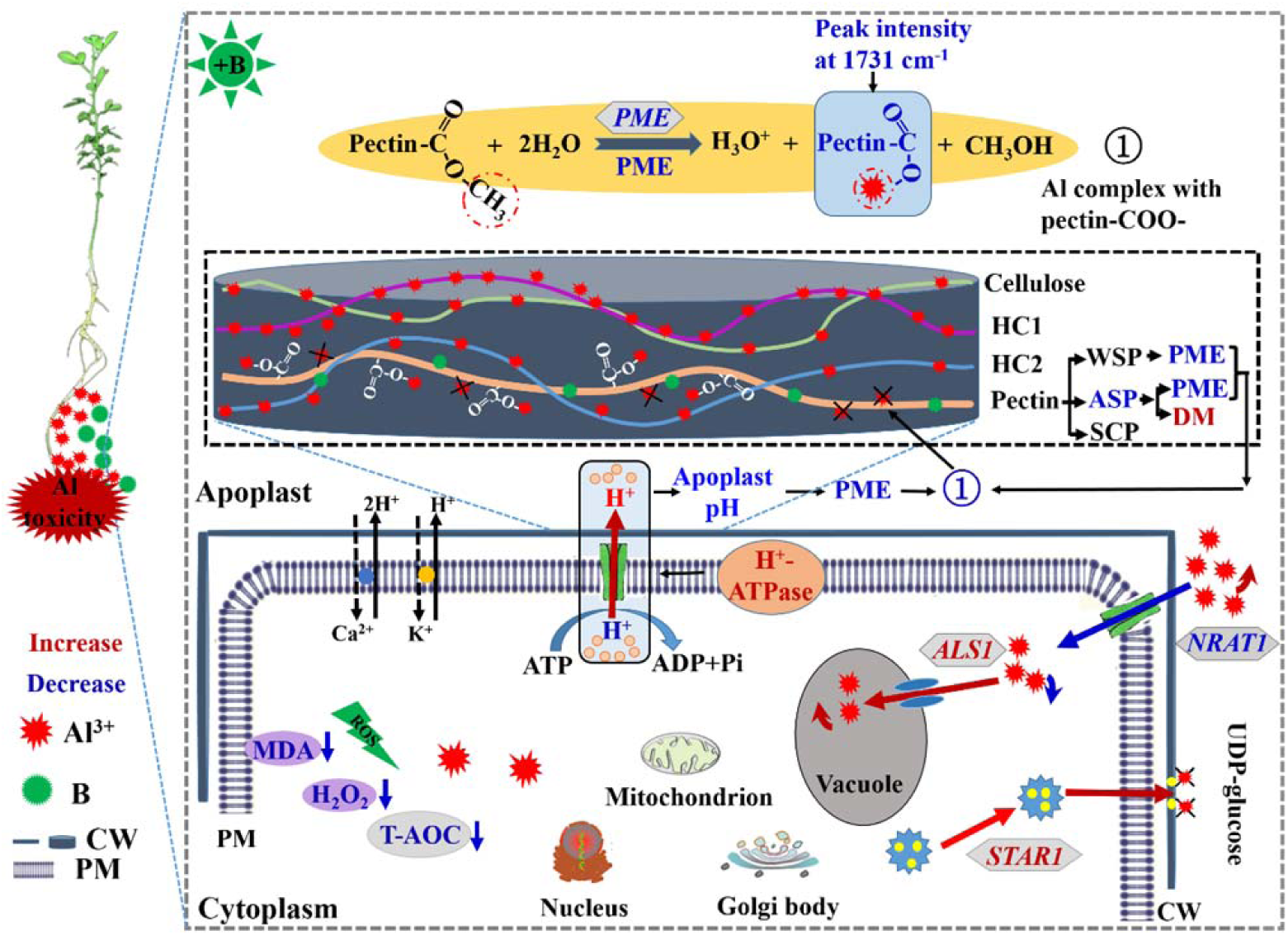
The schematic illustration of a proposed model explains the mechanism of H^+^ transport, pectin methylation, and Al transport and accumulation in root under Al stress. Boron application enhanced H^+^-ATPase activity and promoting H^+^ efflux, which reduced apoplast pH and decreases PME activity, thus decreased Al binding in CW (mainly pectin fractions) and reduced Al toxicity. CW: cell wall; HC1: hemicellulose 1; HC2: hemicellulose 2; PM: plasma membrane; PME: pectin methylesterase; **×**: indicate reduced Al binding in CW.

## Acknowledgments

This work was supported by the National Natural Science Foundation of China (Grant numbers: 41271320) and the Fundamental Research Funds for the Central Universities (Grant numbers: 2017PY055).

---

**Table S1.**
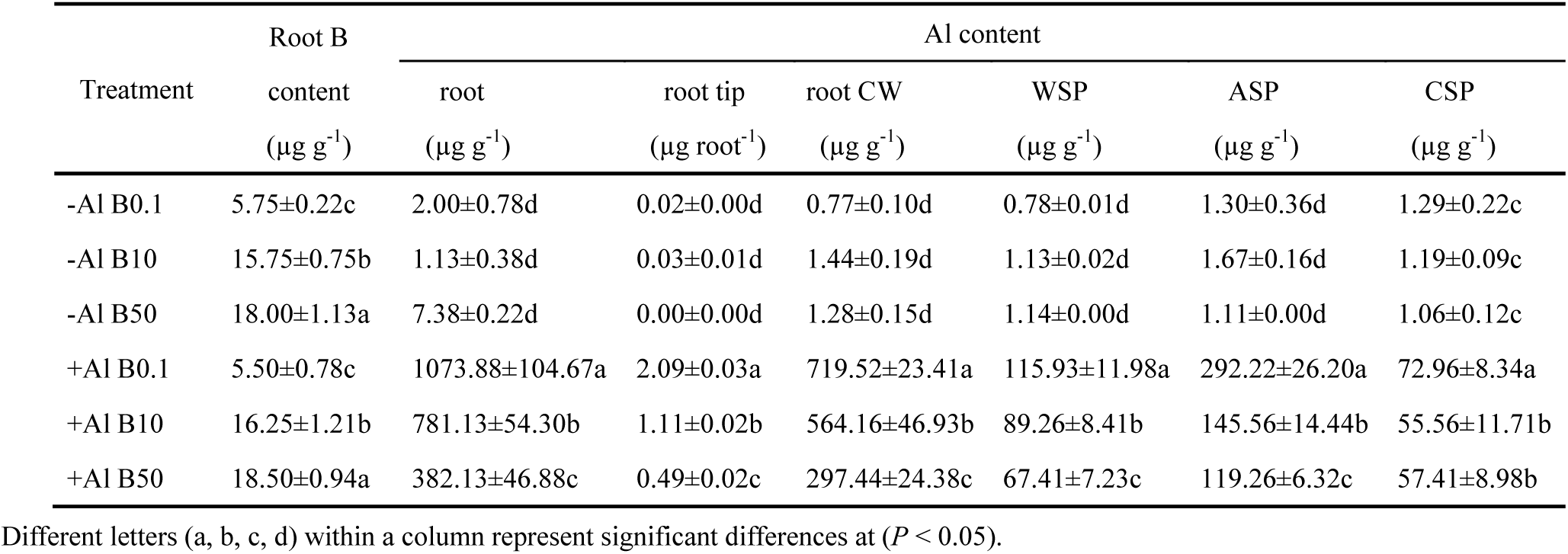
Effects of B on the B and Al contents of trifoliate orange rootstock under Al stress.

| Treatment | Root B | Al content |  |  |  |  |  |
| --- | --- | --- | --- | --- | --- | --- | --- |
|  | content | root | root tip | root CW | WSP | ASP | CSP |
| | ($\mu\text{g g}^{-1}$) | ($\mu\text{g g}^{-1}$) | ($\mu\text{g root}^{-1}$) | ($\mu\text{g g}^{-1}$) | ($\mu\text{g g}^{-1}$) | ($\mu\text{g g}^{-1}$) | ($\mu\text{g g}^{-1}$) |
| -Al B0.1 | 5.75±0.22c | 2.00±0.78d | 0.02±0.00d | 0.77±0.10d | 0.78±0.01d | 1.30±0.36d | 1.29±0.22c |
| -Al B10 | 15.75±0.75b | 1.13±0.38d | 0.03±0.01d | 1.44±0.19d | 1.13±0.02d | 1.67±0.16d | 1.19±0.09c |
| -Al B50 | 18.00±1.13a | 7.38±0.22d | 0.00±0.00d | 1.28±0.15d | 1.14±0.00d | 1.11±0.00d | 1.06±0.12c |
| +Al B0.1 | 5.50±0.78c | 1073.88±104.67a | 2.09±0.03a | 719.52±23.41a | 115.93±11.98a | 292.22±26.20a | 72.96±8.34a |
| +Al B10 | 16.25±1.21b | 781.13±54.30b | 1.11±0.02b | 564.16±46.93b | 89.26±8.41b | 145.56±14.44b | 55.56±11.71b |
| +Al B50 | 18.50±0.94a | 382.13±46.88c | 0.49±0.02c | 297.44±24.38c | 67.41±7.23c | 119.26±6.32c | 57.41±8.98b |

Different letters (a, b, c, d) within a column represent significant differences at ( $P < 0.05$ ).

**Table S2.**
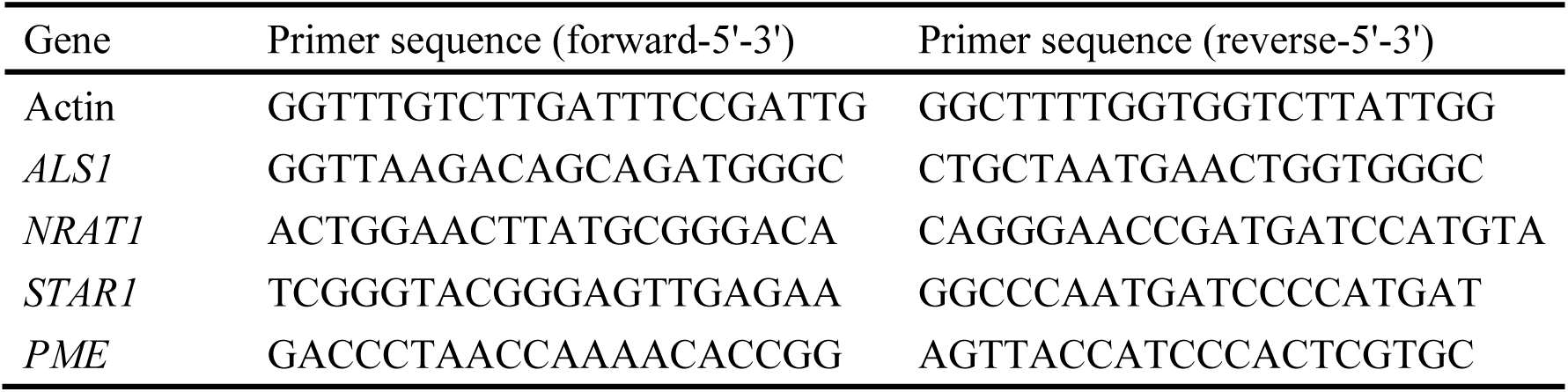
Primers of genes used in this study.

**Table S3.**
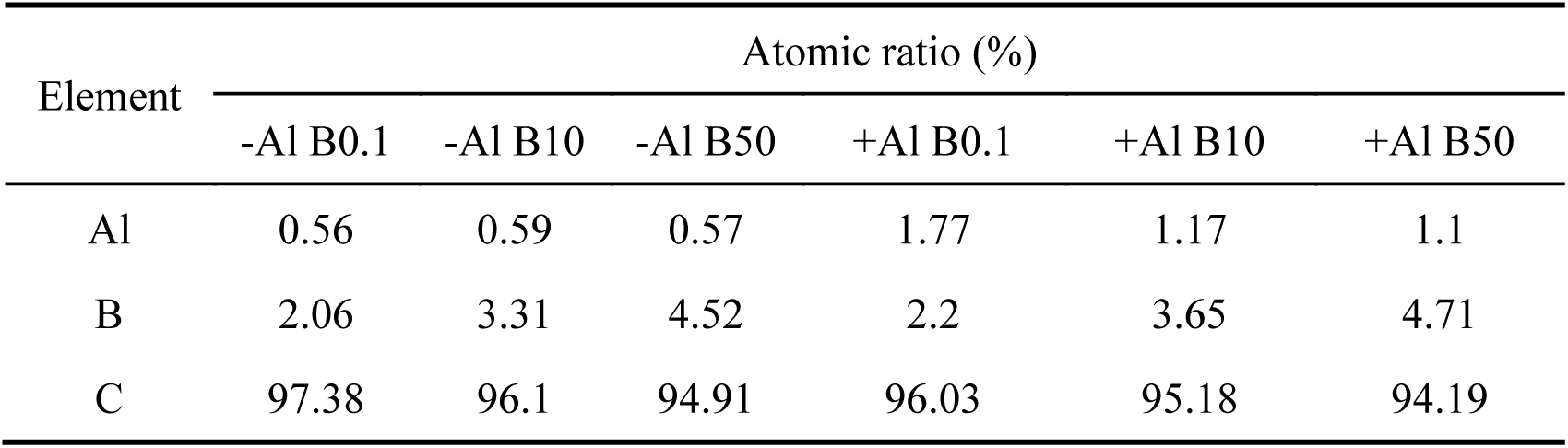
Effects of B on the atomic ratios of carbon, B and Al in root CW by XPS analysis under Al stress.

| Element | Atomic ratio (%) |  |  |  |  |  |
| --- | --- | --- | --- | --- | --- | --- |
|  | -Al B0.1 | -Al B10 | -Al B50 | +Al B0.1 | +Al B10 | +Al B50 |
| Al | 0.56 | 0.59 | 0.57 | 1.77 | 1.17 | 1.1 |
| B | 2.06 | 3.31 | 4.52 | 2.2 | 3.65 | 4.71 |
| C | 97.38 | 96.1 | 94.91 | 96.03 | 95.18 | 94.19 |

**Fig. S1.**
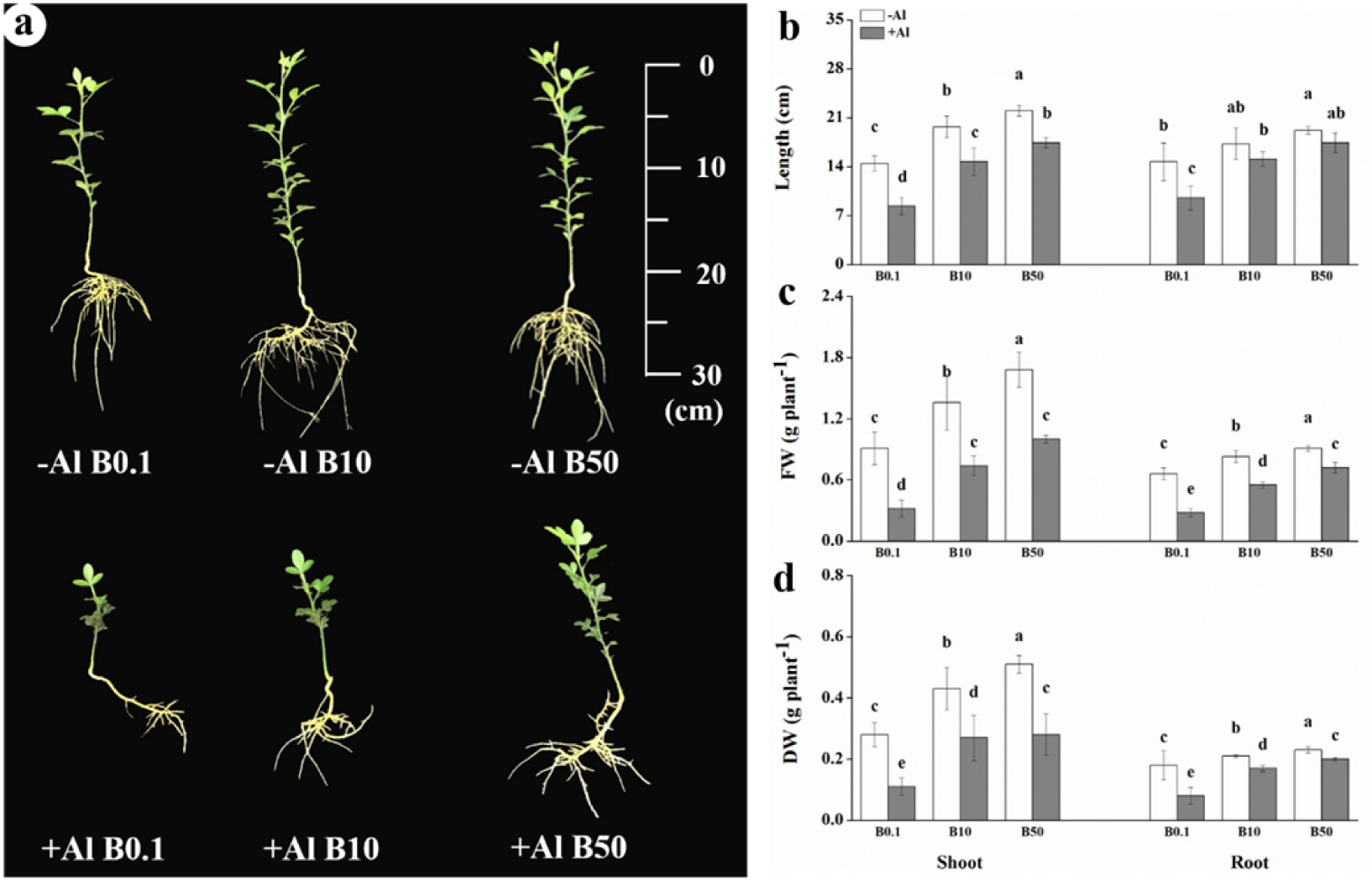
Effects of B on the growth of trifoliate orange rootstock under Al toxicity. The experimental treatments consisted of three B levels (0, 10, and 50 µM as H_3_BO_3_), and two Al levels (0 and 300 µM). Bars are means of three replicates ± SD. Different letters (a, b, c, d) in each sub-figure represent significant differences at (*P* < 0.05).

**Fig. S2.**
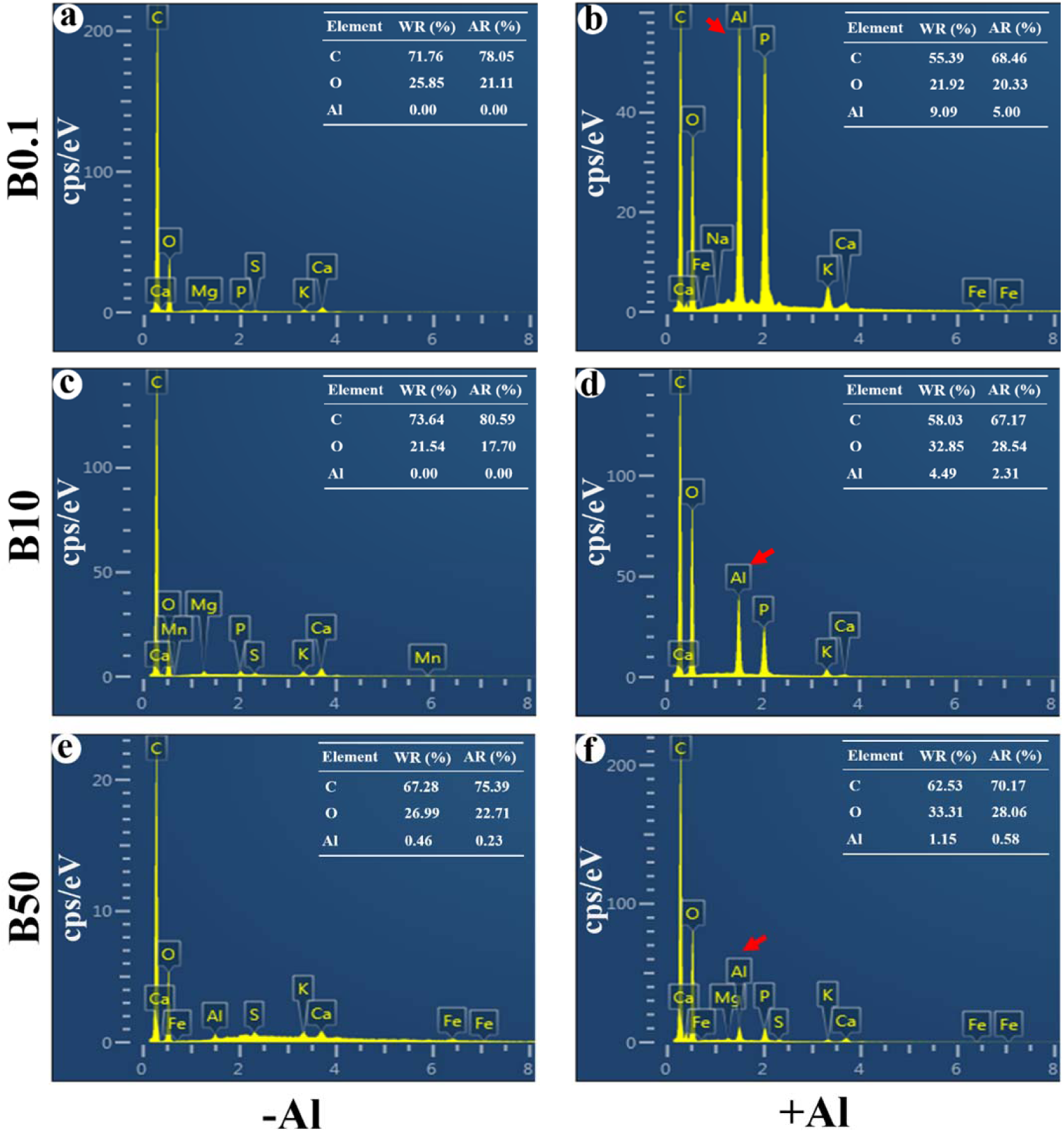
Effects of B on the energy spectrum of Al in in trifoliate orange rootstock root CW under Al stress is reflected by SEM-EDS. Fig-a: Plants treated with 0 µM Al and 0.1 µM B (–Al B0.1); Fig-b: Plants treated with 0 µM Al and 10 µM B (–Al B10); Fig-c: Plants treated with 0 µM Al and 50 µM B (–Al B50); Fig-d: Plants treated with 300 µM Al and 0.1 µM B (+Al B0.1); Fig-e: Plants treated with 300 µM Al and 10 µM B (+Al B10); Fig-f: Plants treated with 300 µM Al and 50 µM B (+Al B50); WR: weight ratio; AR: atomic ratio.

**Fig. S3.**
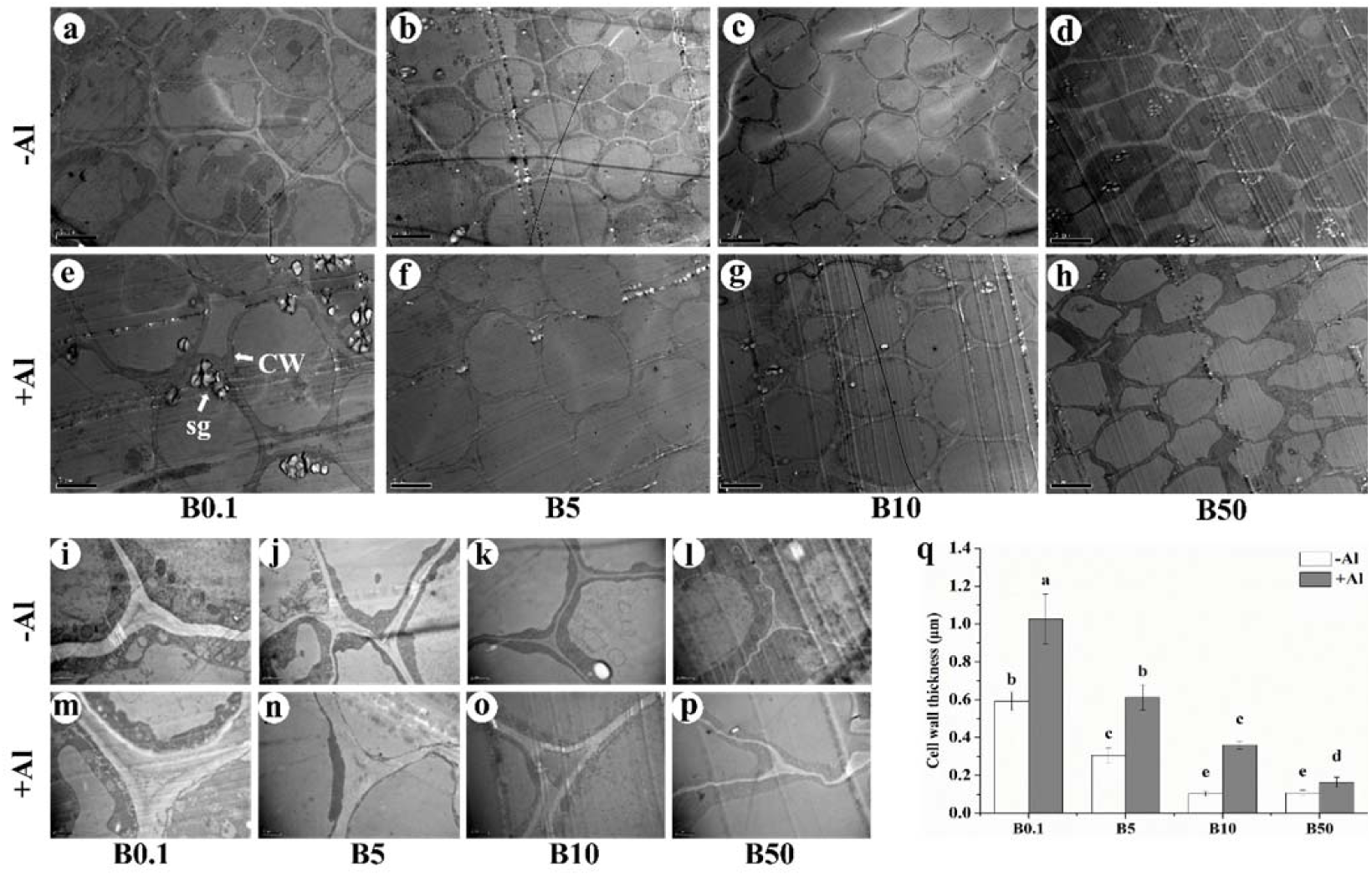
Transmission electron microscope (TEM) of roots of trifoliate orange rootstock that was exposed to B and Al treatments. TEM transverse root section shows the regularity of cell arrangement and the thickness of CW. Additional labelled features of the cell are starch (sg). The experimental treatments consisted of four B levels (0, 5, 10, and 50 µM as H_3_BO_3_), and two Al levels (0 and 300 µM). a-h: Bar = 5 µM; i-p: Bar = 1 µM; Fig-g: CW thickness.

**Fig. S4.**
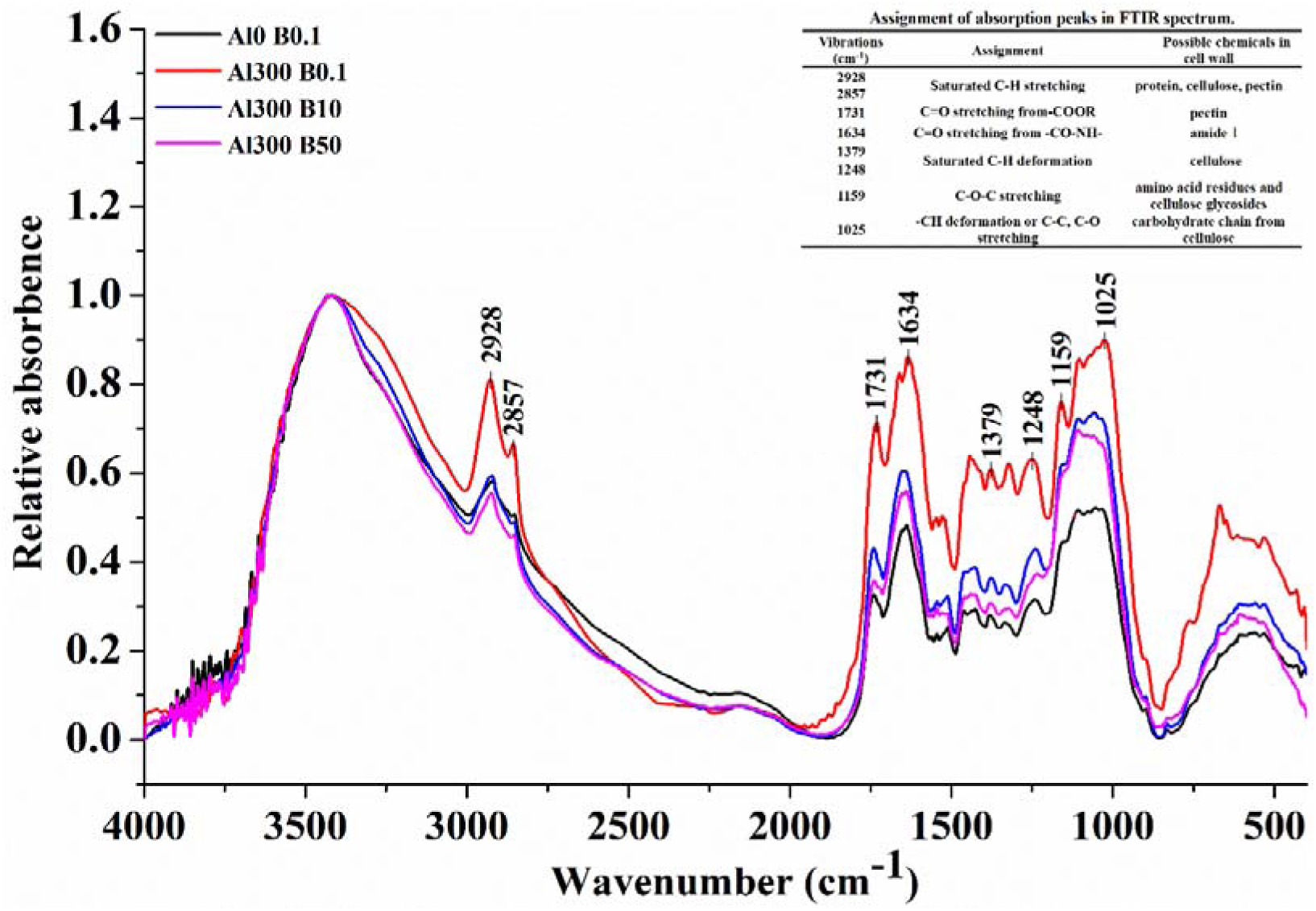
Effects of different B concentrations on the CW composition of trifoliate orange rootstock under Al stress is reflected in FTIR spectra (4000-400 cm^-1^).

**Fig. S5.**
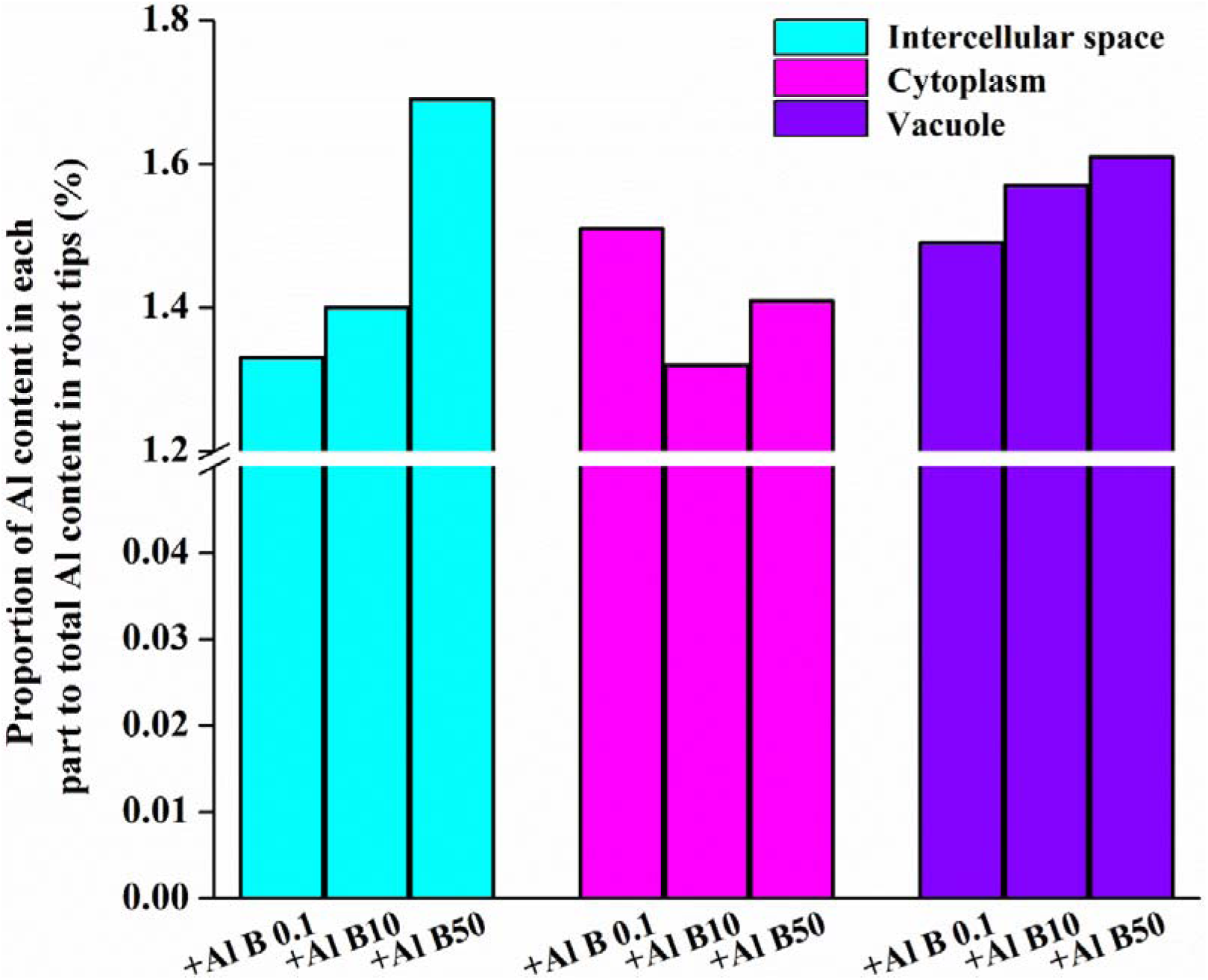
Effects of B on the proportion of Al content in each part to total Al content in root tips under Al toxicity.

**Fig. S6.**
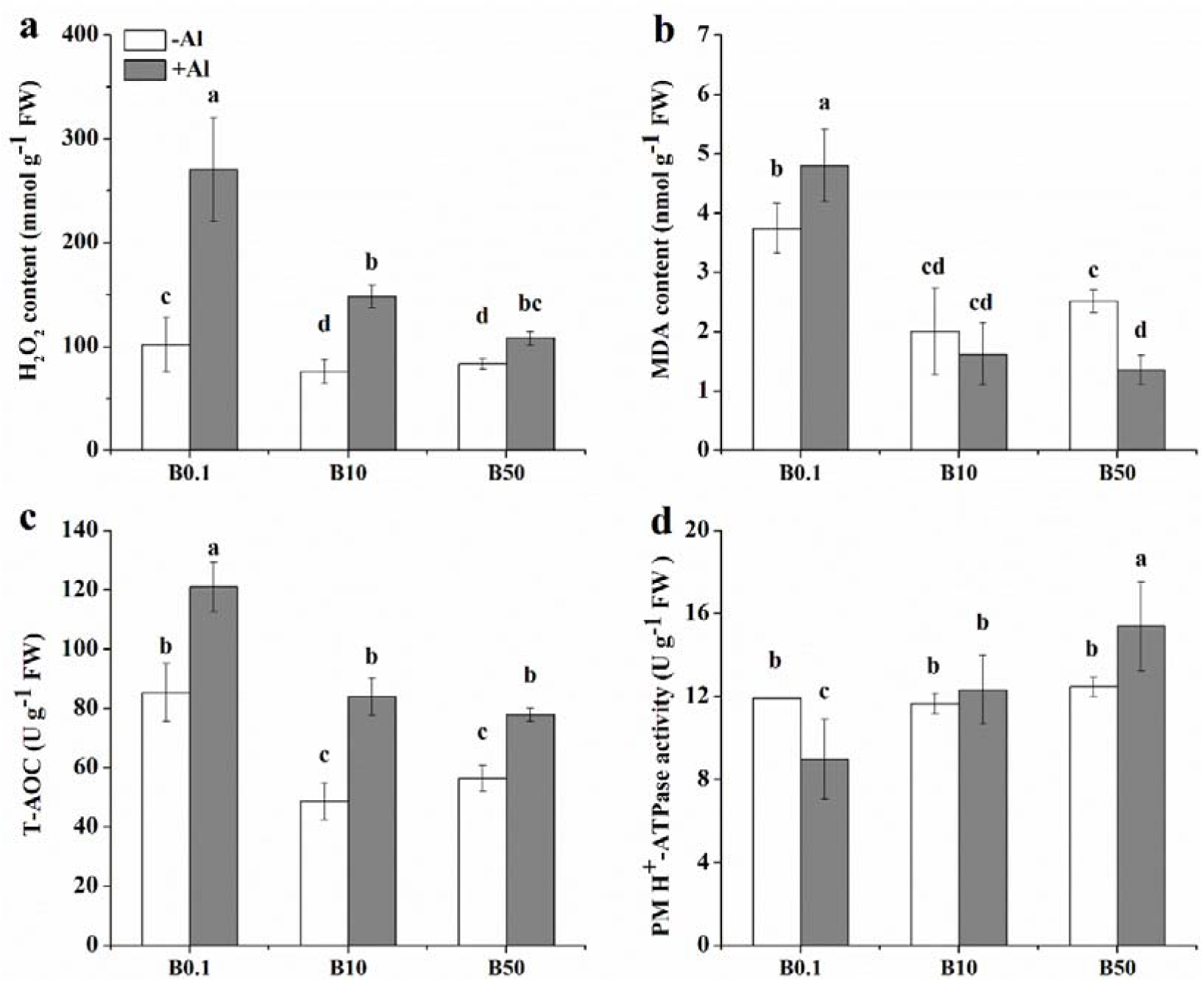
Effects of B on (a) H_2_O_2_, (b) MDA contents, (c) T-AOC, and (d) PM H^+^-ATPase activities in roots under Al toxicity. Bars are means of three replicates ± SD. Different letters (a, b, c, d) in each sub-figure represent significant differences at (*P* < 0.05).

